# Condensate-Like Organization in Respiratory Aerosols Modulates the Dynamics of an Airborne Virus

**DOI:** 10.64898/2026.04.30.721971

**Authors:** Nicholas A. Wauer, Carla Calvó-Tusell, Abigail C. Dommer, Lorenzo Casalino, Fiona L. Kearns, Marcelo Caparotta, Mia A. Rosenfeld, Clare K. Morris, Rommie E Amaro

## Abstract

The molecular behavior of viruses within respiratory aerosols plays a critical role in airborne disease transmission yet remains largely inaccessible to experimental characterization. Here, we use a billion-atom all-atom molecular dynamics simulation of a virus-laden respiratory aerosol to uncover how respiratory proteins, lipids, ions, and water collectively assemble around SARS-CoV-2, giving rise to structured microenvironments that influence viral stability and spike dynamics. We find that respiratory components rapidly evolve into heterogeneous networks characterized by protein-rich aggregates, patchy lipid assemblies, and spatially structured ion and water dynamics. These features create distinct microenvironments that constrain molecular transport and stabilize regions surrounding the virion. Within this crowded aerosol context, we observe sustained and selective interactions between aerosol components and the viral spike protein, including preferential recruitment of surfactant lipids and persistent coordination by divalent cations. These interactions modulate spike conformational dynamics, enhancing domain breathing motions and flexibility at key hinge regions while preserving a stable membrane anchor. Together, these observations reveal a condensate-like physical regime in which multivalent aerosol components coalesce into a soft, heterogeneous matrix that selectively modulates viral protein dynamics under extreme crowding. By framing virus-laden respiratory aerosols within this physical context, this work establishes an *in situ* molecular framework for understanding how aerosols influence viral persistence and offers a platform for exploring mechanisms relevant to airborne disease transmission and mitigation strategies.

**TOC Graphic:** 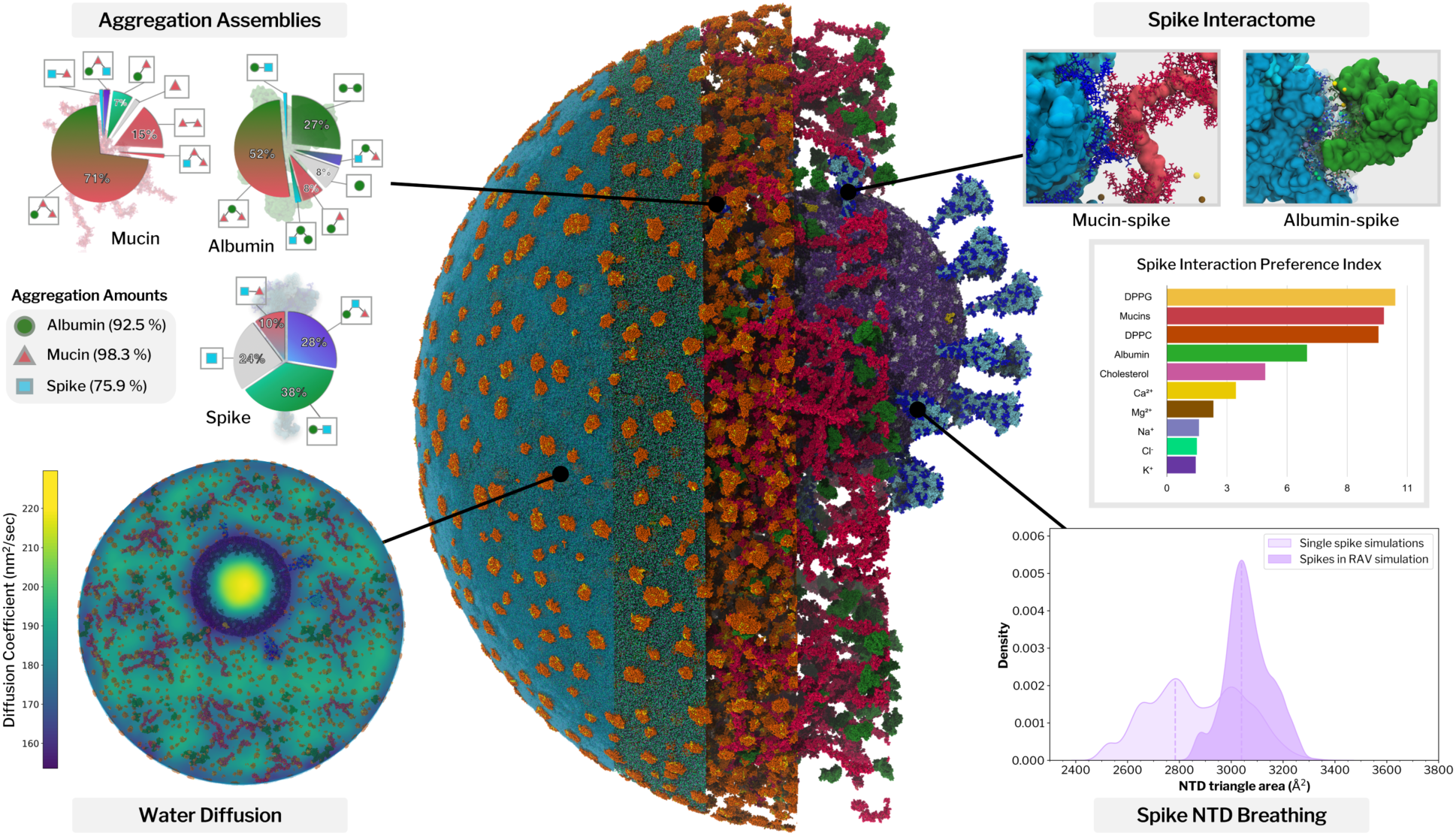

**Synopsis:** Respiratory aerosols exhibit condensate-like physical properties that govern the evolution of the particle and modulate the behavior of airborne SARS-CoV-2.

## 1. Introduction

The recent COVID-19 pandemic, triggered by the emergence of SARS-CoV-2, significantly renewed global interest in understanding the role of aerosols in the spread of disease. Respiratory aerosols (RAs) in particular, emitted while breathing, speaking, and coughing, are efficient carriers of pathogens and are the primary mode of transmission for many viruses^1^. Although RAs can accumulate in closed spaces for minutes to hours^2^, virus infectivity decays due to equilibration of the carrier aerosol with the surrounding environment^3^, which can expose the virion to stressors such as UV-radiation, acidification, and desiccation^4^. Numerous recent studies have shed additional light on how the environment impacts viral infectivity^5–11^, but the specific molecular mechanisms governing virus survival remain largely unknown.

Relative humidity (RH) and the composition of the surrounding gases play a dominant role in virus deactivation. However, emerging evidence suggests that the chemical composition of RAs, most notably the presence of mucins, could mitigate atmospheric damage^12,13^. RAs are derived from the fluid lining the respiratory tract, which is composed of salts, various biological macromolecules (e.g., proteins, lipids, and mucins), and dissolved gases (including high amounts of CO_2_). Upon exposure to ambient air, the rapid loss of CO_2_ coupled with solutes concentrating through particle dehydration can spike aerosol pH, while the dissolution of acidic atmospheric gases such as HNO_3_ into the aerosol reverses the effect^14–16^. As water evaporates in response to changing RH, respiratory mucins, one of the most important glycoproteins in lung fluid, are hypothesized to protect co-aerosolized viruses by preserving them in a semisolid, viscoelastic phase state^12,13,17^. Such a state could hinder the diffusion of solutes, slow water evaporation, and prevent exposure of the virus to deleterious microenvironments such as extremes in salt concentration or pH.^18^

The dynamic evolution of biomolecular structure within aerosol microenvironments remains largely hidden from experimental view; RAs are particularly difficult to collect, characterize, and isolate intact^18^, and common single-particle imaging techniques such as fluorescence microscopy and scanning electron microscopy (SEM) are severely limited in spatiotemporal resolution. However, molecular dynamics (MD) simulations are able to resolve biomolecular systems at molecular and atomistic detail. MD simulations are increasingly employed as a computational microscope to interrogate the mesoscale, most recently exemplified by simulations of a minimal cell^19^, mitochondrial cristae^20^, entire genomes and chromosomes^21,22^, viruses^23–26^, bacterial cytoplasm^27^ and bioaerosols^25,28^. The improved capacity to conduct large-scale, biologically complex simulations paves the way towards understanding how viruses such as SARS-CoV-2 behave *in situ*, forging a path to visualize the molecular interactions and dynamics that shape viral persistence in flight^29^.

Here, we report an all-atom MD simulation of a virus as it exists in the air: a “respiratory aerosol virus” (RAV)—a 270 nm-diameter, billion-atom particle that encapsulates a fully-glycosylated Delta SARS-CoV-2 virion (B.1.617.2 lineage) in a chemically realistic respiratory aerosol. The particle matrix contains the principal components of respiratory fluid—lung surfactant lipids, multiple mucin subtypes, human serum albumin, and physiologically relevant ions—yielding one of the most chemically comprehensive atomistic models of an infectious aerosol to date. Enabled by optimized high performance computing (HPC) workflows, we generated 505.8 ns of production dynamics for this billion-atom system. Through systematic analysis of aerosol organization and virus–environment interactions, we examine how multicomponent protein networks emerge within the particle and how these networks shape the local microenvironment surrounding the virion. We further assess how interactions with respiratory components influence spike conformational behavior under conditions of extreme molecular crowding. Together, these simulations provide a molecular-scale perspective on the physical organization of virus-laden respiratory aerosols and establish a framework for investigating how aerosol composition and internal structure may influence viral stability and dynamics during airborne transport.

## 2. Results and Discussion

### 2.1. System Construction of the Respiratory Aerosol Virus

Applying an integrative modeling approach, we combined an array of experimental datasets to construct a comprehensive representation of SARS-CoV-2 within a respiratory aerosol particle. The Delta SARS-CoV-2 model was informed by cryo-EM, electron tomography, and mass spectrometry, which describe global virion morphology, protein and lipid distributions, and glycosylation patterns^30–32^. The composition of the solute within the RA matrix was adapted from a surrogate lung fluid formula commonly used in experimental studies, the major components of which are DPPC, DPPG, cholesterol, human serum albumin, and mucins^33–35^. The preparation and characterization of the respiratory mucin models used in this work are described in Kearns *et al.*^36^. Respiratory mucins MUC5B and MUC5AC are secreted throughout the respiratory tract and form gels upon chelation with Ca^2+ 37,38^ which we hypothesize could play an integral role in controlling aerosol phase and providing viral protection.

Given that aerosols lose water as they equilibrate with the external environment, the final solute concentrations are elevated compared to the initial concentrations of the lung fluid surrogate. The concentration factor was estimated based on experimental observations of cough aerosol hygroscopicity^40^. It is important to note that the RAV composition is a best estimate, as the actual composition and concentration of matter in RAs is known to be highly variable. Multiple studies show that mass transfer across biological interfaces (such as the sea surface microlayer) can be highly selective due to interfacial enrichment of hydrophobic matter^41–45^, and bioaerosols originating from the same source can exhibit particle-to-particle differences^46–48^. In addition, RA components are observed to vary intrinsically from person to person, and are further modulated by factors such as generation mechanism, disease state, physical fitness, and even time of day^49–53^. We further note that the simulated composition corresponds to an aerosol near the end of the initial rapid drying phase at moderate RH. Moreover, our simulations do not account for reactive atmospheric processing (e.g., aging) or dynamic pH evolution. The final RAV composition approximately matches a 6-fold concentration of the bulk surrogate fluid composition corresponding to a reduction in radial size by almost 50% (**Figure S1-2** and **Table S1**) and represents one relevant state of an externally mixed and heterogeneous population (i.e., the present model captures a protein-rich regime representative of moderate RH cough aerosols). The composite model RAV was then initialized with solute components randomly distributed throughout a sphere, an internal volume was carved out where the virus model was placed, and then the whole particle was surrounded by a vacuum in a cubic box (**Figure 1a and Figure S3, SI Methods**).

**Figure 1.**
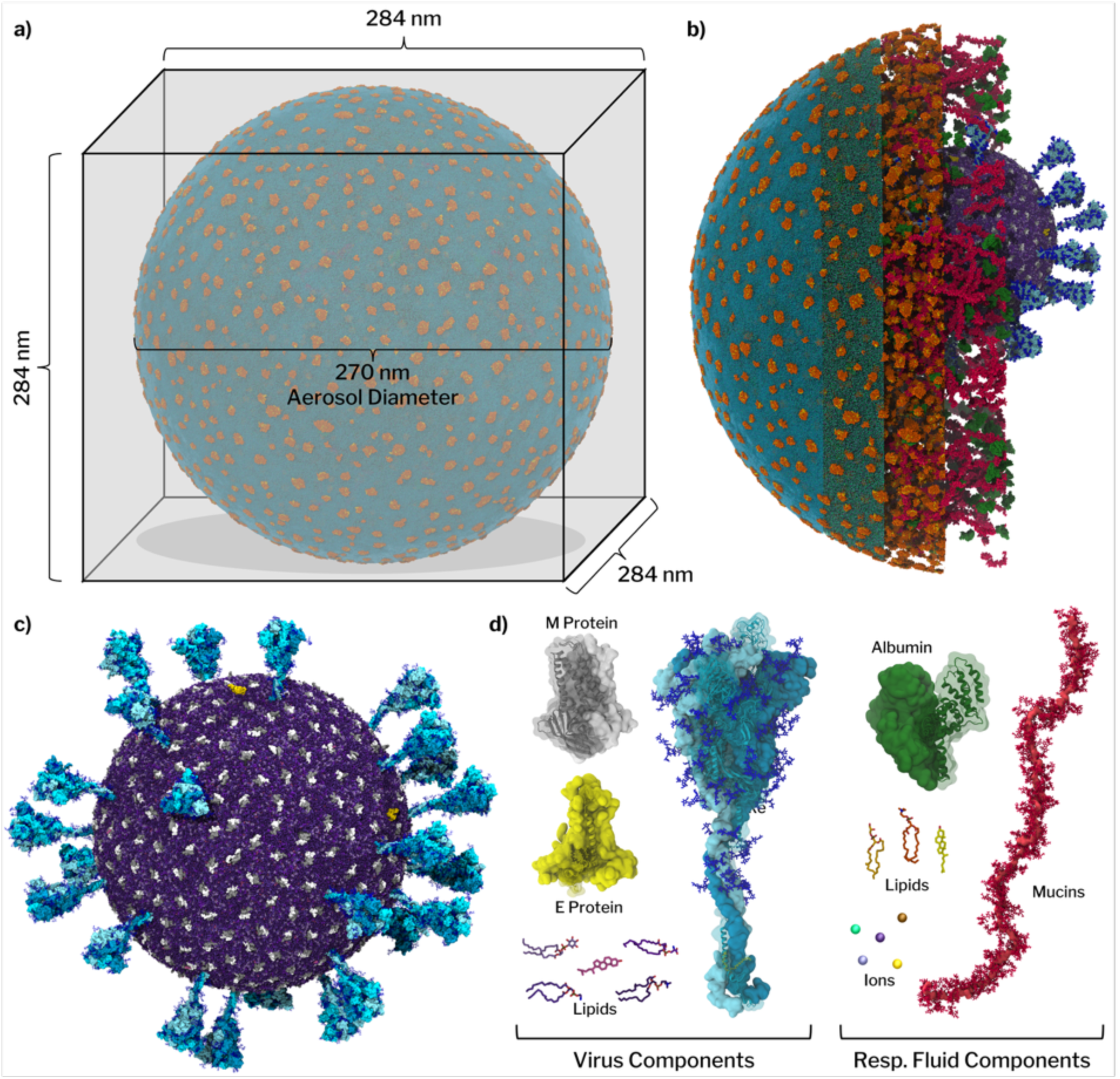
Schematic representation of the respiratory aerosol virus (RAV) system. (a) Illustration of the simulation box where the aerosol is surrounded by vacuum. (b) A cross-sectional rendering of the final simulation snapshot illustrates the various levels of complexity of the RAV particle composition. From right to left, this sliced figure starts showing the SARS-CoV-2 virion, then also including mucins and albumins, then lipids, then ions and finally the complete model including water molecules, which can also be seen in (a). (c) Full render of the SARS-CoV-2 delta virion present within the aerosol system. (d) On the left, the components of the embedded SARS-CoV-2 Delta virion are shown. The virion is formed by 29 spike trimers, 356 M-protein dimers, and 4 E-protein pentamers, together with the viral envelope lipids (POPC, POPE, POPI, POPS, cholesterol). On the right, the components of the respiratory fluid are shown. Respiratory fluid is composed of mucins, albumins, surfactant lipids (DPPC, DPPG, cholesterol), and inorganic ions (Na⁺, K⁺, Cl⁻, Ca²⁺, Mg²⁺). Full compositional data can be found in the extended methods and Table S1.

### 2.2 Molecular Transport and Spatial Heterogeneity of the Respiratory Aerosol Matrix

We sampled ∼500 ns of all-atom production molecular dynamics of the aerosol in the cubic box, with periodic boundary conditions, to capture early reorganization within the respiratory aerosol. Over this timescale, ∼6000 (0.002%) water molecules evaporated into the vacuum region to establish equilibrium across the gas-liquid interface **(Figure S4)**, which puts the gas phase at an estimated ∼72% RH (see SI Methods). Substantial redistribution and reorganization occurs within the aerosol interior and at the surface, indicating that internal transport, not bulk deformation, dominates early RAV evolution.

To characterize molecular mobility within this crowded environment, we computed self-diffusion coefficients for low–molecular weight aerosol components. Ion diffusion **(Figure S5)** reveals pronounced heterogeneity: monovalent ions (Na⁺, K⁺, Cl⁻) remain relatively mobile and display bulk-like behavior in water-rich regions, whereas divalent ions (Ca²⁺, Mg²⁺) diffuse more slowly due to stronger and longer-lived electrostatic interactions with proteins, glycans, and lipids. Among divalent species, Ca²⁺ is overrepresented in the low-diffusion regime, consistent with its propensity for persistent coordination with higher-order respiratory components **(Figure S5 and Section 2.4)**. Spatial ion distributions further show enrichment of Ca²⁺ and Mg²⁺ near the virion surface, while monovalent ions approach closer to the air–water interface, suggesting preferential sequestration of divalent ions within the crowded interior **(Figure S6)**.

Respiratory lipids (DPPC, DPPG, and cholesterol) remain highly mobile within the aerosol matrix and undergo rapid self-organization **(Figures S7-S10)**. Initially dispersed, these lipids reorganize into micelle-like clusters and surface-associated patches, making lipid mobility a major driver of early RAV morphological evolution (**Figure 2a and Figure S8**). Density-based clustering analysis shows rapid nucleation of many small lipid aggregates, with cluster numbers peaking near 100 ns **(Figure S9)**. Thereafter, clusters gradually merge and grow as free lipids are recruited, leading to larger lipid-rich assemblies. By the end of the simulation, approximately 90% of respiratory lipids participate in clusters, with the population of free lipids decaying exponentially over time.

**Figure 2.**
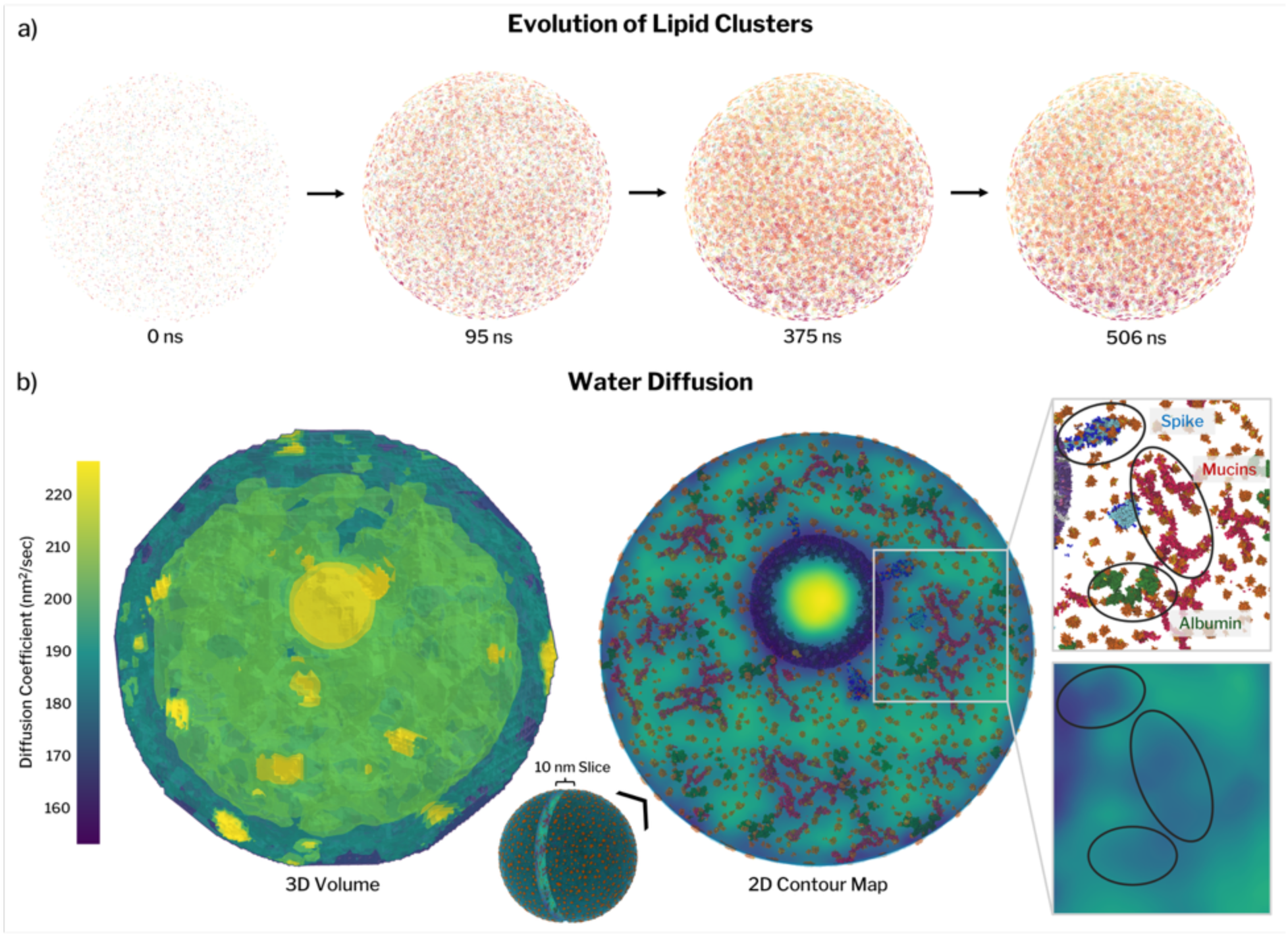
Dynamic morphology and reorganization in the respiratory aerosol virus (RAV). (a) Lipid clustering dynamics. Time evolution of respiratory surfactant lipids (DPPC and DPPG) showing rapid nucleation from an initially dispersed distribution into many small aggregates that progressively merge over the 500 ns trajectory (snapshots at 0, 95, 375, and 506 ns). (b) Spatial heterogeneity of water diffusion. Three-dimensional map of local water diffusion throughout the aerosol volume (left) and a two-dimensional contour map from a 10 nm-thick slice (right), overlaid with the positions of higher-order components (spikes, mucins, and albumins). Local diffusion values were obtained from mean-square displacement–based analysis within spatial voxels (see **Methods, Figure S11, Movie M1**).

Restricting the analysis to lipids located at the air–water interface reveals similar aggregation kinetics. Surface-associated lipid clusters form rapidly, followed by a monotonic decrease in cluster number as cluster size increases **(Figure S9)**. Surface coverage grows quickly during the initial diffusion-dominated phase but slows thereafter, reaching a maximum of ∼9.7% of the particle surface area. Complete surface saturation would require either substantially more lipids than are present or the continued reduction in particle size *via* decreased RH. Even if all surfactant lipids were localized at the interface, coverage would reach only ∼76%. Thus, under these conditions, surfactant lipids form a patchy, discontinuous surface layer rather than a continuous monolayer, a feature unlikely to influence surface transport and accessibility to atmospheric reactants due to the high amount of unobstructed air/water surface area.

This is contrary to prior sea spray aerosol simulations where organic aggregates lead to significant structural and morphological deformations. The RAV model experiences similar rearrangement and aggregation kinetics, but the overall organization remains highly distributed with no dominant lipid structures emerging. The behavior is observed for the surface structure which in the RAV exhibits disconnected lipid patches while the SSA models reach a saturated or nearly saturated particle surface. Most of these differences are likely due to compositional differences, mainly the difference in overall organic content, with the SSA models having an organic mass percent of ∼50% whereas the RAV is only ∼10%. Additionally, the scale of these particles leads to these differential outcomes as the SSA particles are∼40nm in diameter and the RAV is ∼270nm in diameter. These systems were simulated for a similar timescale but the drastic difference in size and organic mass fraction means many of the large-scale morphological rearrangements seen in the SSA are not observed in the RAV due to the diffusion limited movements of lipid clusters and the overall more dispersed organization of the organic material.

To integrate these observations into a spatially resolved picture of transport, we quantified local water diffusion throughout the aerosol using voxel-based mean-square displacement analysis. Three-dimensional diffusion maps reveal strong spatial heterogeneity (**Figure 2b, Figure S11, Movie M1**). Water mobility is reduced near the air–water interface and in crowded regions surrounding the virion, while the virion interior—containing only water and NaCl—exhibits higher, bulk-like diffusion values, serving as an internal control. Localized regions of elevated diffusion at the particle surface coincide with sites of water evaporation.

Mapping diffusion coefficients onto aerosol composition shows that regions enriched in spike proteins, mucins, and albumins exhibit markedly reduced water mobility (**Figure 2b**). The cage-like patterns observed in diffusion maps reflect extended mucin–albumin networks, with occasional connections to the spike protein. Quantitative correlation analysis (Pearson coefficient, **Figure S12, Table S2**) confirms that water diffusion correlates positively with solvent and monovalent ion content, and negatively with viral membrane proximity and protein density. Regions of protein aggregation consistently correspond to zones of restricted transport. Such spatially heterogeneous transport and composition-dependent mobility are hallmarks of soft, viscoelastic assemblies formed by multivalent macromolecular interactions ^53,54^. These features indicate that the respiratory aerosol behaves as a dynamically structured matrix rather than a homogeneous solution.

### 2.3. Emergent Mesoscale Structure: Mucin and Protein Networks in the Respiratory Aerosol

A central question is whether the respiratory proteins and mucins merely coexist with the virion or whether they actively reorganize to form structured interaction environments around it. Such reorganization could influence local crowding, transport, and the physicochemical context experienced by viral surface proteins. To address this, we analyzed protein–protein aggregation patterns within the aerosol using a network-based framework in which proteins are represented as nodes connected when they lie within a defined spatial distance of one another. Varying this distance cutoff, i.e., the maximum separation at which two proteins are considered part of the same aggregate, allows aggregation to be probed across increasing length scales, from local contacts to extended mesoscale assemblies.

A distance-based graph network analysis of protein aggregation reveals that mucin and albumin readily cluster under moderate distance cutoffs (e.g., 100–200 Å, **Figure 3 and Figure S13-S16**). These interactions can lead to the formation of sizable protein-rich microenvironments, particularly in the interior regions near the virion or in patches where mucins act as bridges between albumin clusters. In contrast, spike proteins tend to occupy peripheral positions in the aggregation network, consistent with limited participation in the mucin-albumin clusters. At a 200 Å threshold, most macromolecules participate in aggregates (mucins 98.3%, albumin 92.5%, spikes 75.9%; **Figure 3d**), with mucin-centered assemblies dominated by mucin-albumin networks (71%) and mucin–mucin contacts (15%), and albumin assemblies split between mixed mucin-containing clusters (52%) and albumin–albumin clusters (27%). In contrast, spike-containing assemblies are biased toward mixed aggregates with albumin (38% spike–albumin and 28% spike–albumin–mucin), with fewer spike-only (24%) groupings, consistent with more limited integration of membrane-anchored spikes into the mucin–albumin network. The overlap between slow-diffusing nodes and protein-rich aggregates highlights how local crowding (often driven by mucins and albumins) hampers aerosol component mobility.

**Figure 3.**
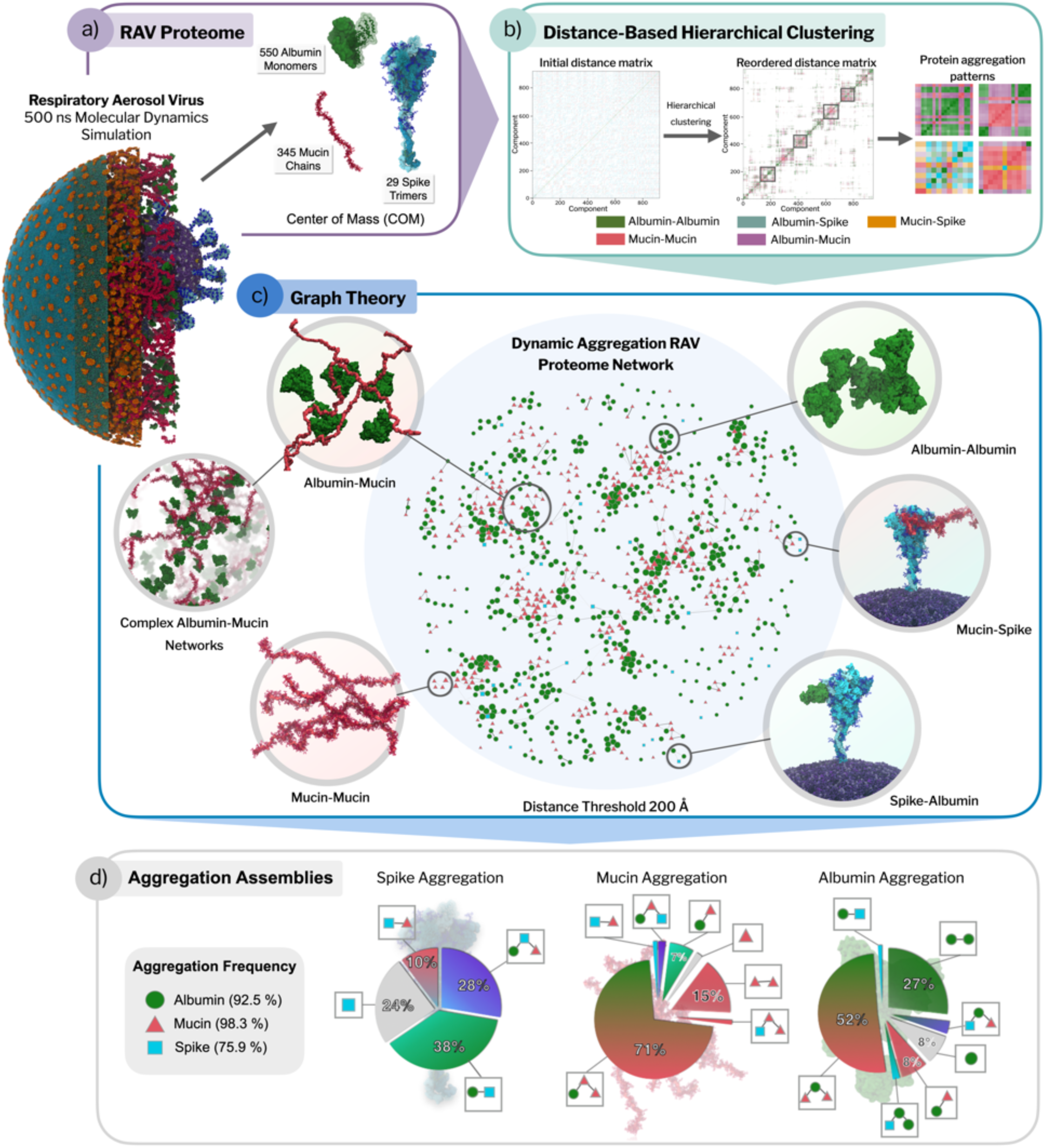
Dynamic macromolecular aggregation in the respiratory aerosol virus. (a) RAV proteome representation. Overview of the macromolecular components analyzed over the 500 ns RAV trajectory, including 550 human serum albumin monomers, 345 mucin chains, and 29 membrane-anchored spike trimers. Proteins are represented by their centers of mass (COM) to enable time-resolved aggregation analysis. (b) Distance-based aggregation workflow. For each trajectory frame, pairwise COM distance matrices were computed and subjected to distance-threshold–based hierarchical clustering to identify protein aggregates and classify their compositions (albumin–albumin, mucin–mucin, albumin–mucin, albumin–spike, and mucin–spike; Methods). Reordered distance matrices highlight emergent aggregation blocks and recurring interaction patterns. (c) Graph-based aggregation network. A representative “proteome network” is shown for a permissive cutoff (200 Å), where nodes correspond to individual proteins (albumin, mucin chains, spike) and edges indicate proximity/association under the distance criterion. (d) Aggregate composition statistics. Summary of aggregation propensity and co-assembly modes across the trajectory. The left panel reports the fraction of each protein type found in aggregates (albumins 92.5%, mucins 98.3%, spikes 75.9%). Pie charts categorize the dominant “assembly types” for each component (spike-, mucin-, and albumin-centered aggregates).

Our analyses reveal that the aerosol proteome does not remain randomly distributed, but instead forms dynamically evolving, heterogeneous local environments characterized by varying degrees of crowding and connectivity. The spontaneous emergence of such multivalent assemblies parallels organizing principles described for biomolecular condensates, in which scaffold-like macromolecules define a dynamic matrix that transiently engages client components^53–56^. Importantly, the emergence of this structured matrix implies that viral components experience distinct microenvironments depending on their local protein context. This motivates a closer examination of how these assemblies interact with and modulate the SARS-CoV-2 spike protein.

### 2.4 Spike Microenvironment and Dynamics in the Respiratory Aerosol

#### 2.4.1. Early Reorganization of the Spike Protein Interactome

Having established that the respiratory fluid environment reorganizes into structured lipid and protein assemblies, we next examine how these emergent microenvironments interact with and modulate the SARS-CoV-2 spike protein. Specifically, we analyze the composition, dynamics, and functional consequences of the spike’s local molecular neighborhood. We characterized the spike interactome (29 spike proteins in total) by tracking the molecules that populate the spike’s first contact shell (6 Å) over time, revealing the progressive reorganization of the local environment via dynamic exchange and selective retention of certain components (**Figure 4a and Figures S17-S18**).

**Figure 4.**
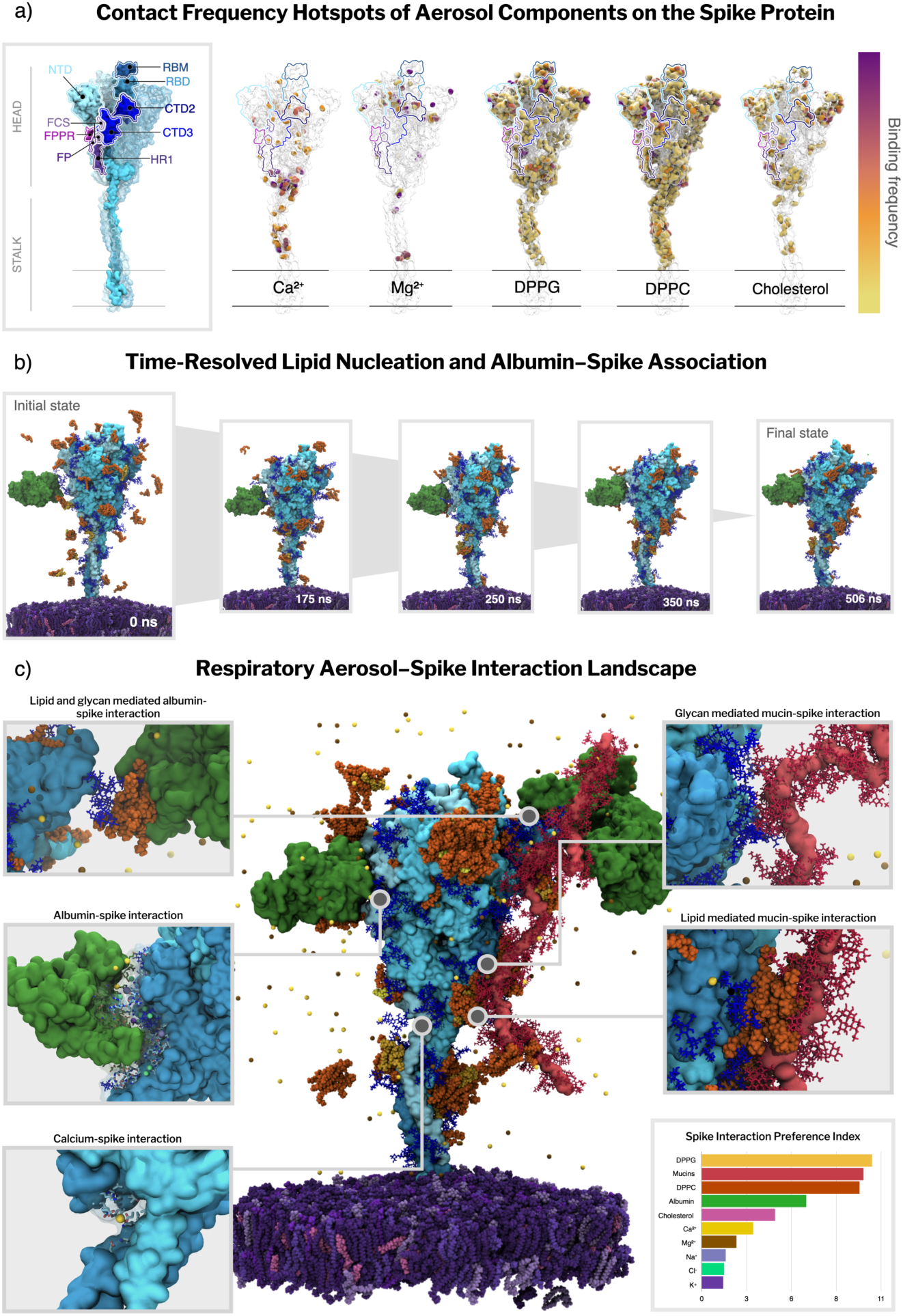
Spike microenvironment in the respiratory aerosol. (a) Contact-frequency hotspot maps on the spike. Residue-level contact frequencies for major aerosol components (DPPG, DPPC, cholesterol, Ca²⁺, Mg²⁺) mapped onto the spike surface. Frequencies were normalized by the number of analyzed frames to obtain a time-averaged contact frequency (ranging from 0, no contact, to 1, contacted in all frames). Plasma color scale indicates no contact (white), low (yellow) and high (purple) contact frequency. (b) Time-resolved lipid nucleation and albumin association. Representative snapshots illustrating progressive accumulation of aerosol surfactant lipids on the spike surface across the 500 ns RAV trajectory (0, 175, 250, 350, and 506 ns). (c) Respiratory aerosol-spike interaction landscape. Representative MD snapshot highlighting a spike embedded in the viral membrane together with representative aerosol components in its vicinity (see **Movie M2**); insets show mechanistically distinct interaction modes observed in the trajectory: lipid- and glycan-mediated albumin–spike contacts, glycan-mediated mucin–spike contacts, lipid-mediated mucin–spike contacts, and site-specific Ca²⁺ coordination at the spike surface and spike interaction preference index. The preference index is shown as a bar plot reporting the relative enrichment of aerosol components within the first interaction shell (6 Å) of the spike. Color scheme: spike protein, cyan/blue (domains as annotated in panel a); viral membrane, purple; mucins, magenta/red; human serum albumin, green; surfactant lipids (DPPC/DPPG/CHOL), orange; glycans, dark blue; ions, yellow/gold (with Ca²⁺ highlighted in panel b inset). See Methods for contact definitions, frequency and preference index calculations.

Over the course of the simulation, the ion composition of the spike’s contact shell experiences a depletion of Na^+^ and Cl^−^ and an enrichment of Ca^2+^ from the initial distribution (**Figure S17**). This suggests that while Na^+^ and Cl^−^ are being outcompeted for binding locations by other components, Ca^2+^ is preferentially binding more than any of the other ions and potentially more than the other solutes. To gain further insight into the preferred hotspots of divalent ions, we mapped their residue-level contact frequencies and visually inspected their positioning over time (**Figure 4a and Figure S18**). Calcium ions exhibit clear site-specific preferences, accumulating significantly at the stalk-head junction (hip), the knee region, and the lower region of the spike head. These binding sites often involve coordination between acidic residues, nearby glycans, and adjacent lipid clusters (**Figure S19**). Persistent calcium occupancy is also observed in internal pockets, and additional ions are localized near N234 and the fusion domain regions implicated in infectivity-enhancing stabilization^57^. These observations support a possible mechanistic role for Ca^2+^ in modulating hinge-like flexion and maintaining local structure under aerosol conditions. Additional calcium ions are seen coordinated at the mucin-spike and albumin-spike interface. Magnesium, though overall less abundant, exhibits persistent binding at similar sites, including the hip region and near the base of the head (**Figure 4a and Figure S18**). Such progressive exchange and selective retention of specific components are characteristic of interaction-driven partitioning in heterogeneous soft-matter assemblies^53,58^, rather than nonspecific adsorption in dilute solution.

Over the first ∼200 ns, transient ionic interactions are progressively lost and the spike microenvironment becomes dominated by longer-lived lipid and protein contacts, prompting us to examine lipid nucleation around the spike. MD simulations show a lipid-nucleation process that progressively covers the spikes in a heterogeneous corona (**Figure 4b, Movie M2**). The two dominant lipid surfactants, DPPC and DPPG, increase three- to four-fold within the MD simulations, reaching above 120 and 15 molecules per spike, respectively, while minor species such as cholesterol also follow a similar trend (**Figure S20**). Interestingly, conformational heterogeneity of spike protein seems to modulate this recruitment: spikes that display an open RBD conformation attract around 30% more DPPC, DPPG, and cholesterol than their closed counterparts, whereas the ion trends are essentially conformation-independent (F**igures S20 and S21**). Respiratory lipids do not adsorb randomly over the spike protein but concentrate into well-defined hotspots near glycans. Contact frequency data show that the highest-occupancy lipid clusters are found at the crown–stem junction at the NTD-stalk interface and along the outer face of the receptor-binding domain (RBD). In both locations, the majority of DPPC/DPPG establish contacts between lipid headgroups and terminal glycan moieties (N165, N234, and N343 glycans), which in turn anchor the surfactant tailgroups against hydrophobic sidechains on the protein scaffold. These glycan-mediated interactions generate a contiguous lipid belt that increases by 30% when the RBD adopts an open conformation, effectively enveloping the exposed RBM **(Figure S20 and Figure S22)**. A second region of lipid binding is located at the base of the stalk; here, lipid clusters interact with the ankle and knee regions. Cholesterol, although less abundant, localizes in the same regions, suggesting a potentially cooperative assembly of mixed lipid microdomains. Together, these results indicate that spike glycans actively template the formation of a surfactant lipid corona that remodels spike exposure to the crowded aerosol microenvironment and can mechanically tune spike flexibility, potentially shielding the protein surface from inactivating agents in the aerosol milieu.

Due to the low diffusion rate of the aerosol proteins (Figure S13), the spike’s protein neighbors remain relatively sparse throughout the simulation. However, when present, albumin and mucin interactions with the spike are notably stable over time. Visual inspection of the trajectory reveals that mucin-spike interactions occur preferentially at the spike head, particularly in the NTD and RBD regions, and involve only the glycans of the mucins, which contact both spike glycans and exposed residues (**Figure 4c**). These interactions are often separated by intercalated lipid clusters that form hydrophobic patches between the mucin glycans and the spike surface. Albumin-spike interactions display greater mechanistic variability: some albumins directly bind the spike via RBD residues and glycans, while others associate indirectly through lipid clusters and persistent ion coordination. Salt bridges between charged residues of the spike and albumin are in some cases further stabilized by coordinated ions and anchored lipids (**Figure 4c**). These spike-aerosol protein interfaces demonstrate that, while not frequently observed in our simulations, likely due to the limited mucin concentration and the complexity of the aerosol system, albumin and mucin contacts with the spike are highly structured and long-lived, shaped by a network of protein-glycan, glycan-lipid, and ion-mediated interactions.

Together, these data reveal a significant dynamic reorganization of the spike’s molecular neighborhood along the MD trajectory. Initial ionic interactions rapidly reorganize and spatially resolved lipid clusters form. Mucin and albumin engage the spike in persistent interactions, often stabilized by lipid and ion scaffolds. This remodeling likely modulates the exposure, flexibility, and structural stability of the spike under crowded conditions present in the aerosol phase.

#### 2.4.2. Reorganized Chemistry of the Spike Microenvironment

To distinguish genuine molecular affinity for the spike protein from mere environmental abundance, we computed a preference index that normalizes the time-averaged occupancy of each component within 6 Å of the spike relative to its bulk concentration (**Figure 4c and Figure S23**). This normalization “corrects” for abundance-driven bias, where highly prevalent species dominate raw contact counts simply because they are more available in the aerosol, not necessarily because they are preferentially retained at the spike surface (**Methods**). Within the spike interactome, mucins and the two dominant surfactant lipids, DPPC and DPPG, exhibit the strongest relative preferences for the spike surface. Together with the sustained build-up of DPPC and DPPG seen in **Section 2.4.1**, these values indicate that the spike does not indiscriminately interact with surfactant lipids; rather, it selectively nucleates lipids whose head groups can hydrogen-bond to its glycan shield or acidic residues. Albumin also shows moderate preference, often forming stable interactions with the spike. Cholesterol is also moderately enriched, frequently co-localizing with phospholipid clusters in glycan-rich surface regions. While most ions are depleted—quantitatively supporting a largely nonspecific electrostatic background—Ca²⁺ retains the highest preference of any ion for the spike surface. Consistent with the diffusion analysis of **Section 2.2**, Ca^2+^’s reduced mobility translates into longer residence times at the spike surface.

We also calculated preference indices centered on mucins and albumin, revealing how these major biomacromolecules interact with the spike and other molecules in the respiratory fluid (**Figure S24**). Albumin shows a marked self-association and a strong preference toward mucins, but only a mild attraction to spikes, supporting the idea that albumin tends to embed in the protein and mucin network rather than interacting strongly with the virion directly. Similarly, mucins exhibit a strong preference for interacting with other mucins, but only a weak affinity for the spike protein. This discrepancy with the spike’s preference index suggests that despite the spike having a strong preferential interaction with mucins, mucin-mucin interactions are so much stronger that the spike interactions are outcompeted. Together, these results support a mesh-like mucin network that surrounds the virion without strongly adhering to it over the simulated timescale. Meanwhile, lipids and ions are partially excluded from these albumin and mucin protein-protein interaction zones, highlighting the chemical compartmentalization and preference toward the spike.

The preference landscape reveals a coordinated reorganization of molecules around the virion and spike protein. Monovalent ions are shed early, while surfactant lipids are actively recruited to the spike surface. Mucins and albumin co-assemble into an associated proteinaceous shell that does not tightly adhere to the virion but impacts the surrounding microenvironment by modulating diffusion and coalescing organics around the virus. As the particle continues to age and shrink in size, this effect is expected to become even more prominent and shield the virus’s microenvironment from potential deactivating agents such as pH changes or dissolved atmospheric gases. Notably, Ca²⁺ emerges as an outlier among ions: despite overall ion depletion, it remains the most preferentially retained at the spike surface, consistent with its reduced mobility and longer residence times. Such selective, concentration-normalized enrichment is characteristic of interaction-driven partitioning in condensate-like assemblies, where local composition is governed by specific molecular affinities rather than bulk abundance ^53,58^. The preference index thus provides a quantitative framework to evaluate how changes in particle composition, humidity or spike mutations can reshape the virion’s immediate environment.

#### 2.4.3 Aerosol-Induced Modulation of Spike Conformational Dynamics

To explore how the reorganized aerosol microenvironment modulates spike conformational dynamics, we analyzed all 29 spike proteins over the final 352 ns of production dynamics (∼10.2 µs cumulative sampling). Molecular crowding is often assumed to restrict protein motion under conditions of hard steric confinement. To test whether this holds in a RA, we quantified the three principal hinge angles that govern spike orientation: the hip (head-to-upper-stalk), knee (upper- to lower-stalk), and ankle (lower-stalk to membrane normal) (**Figure 5).** Compared to an in-solution single-spike reference ensemble (10.2 µs of wild-type simulations; **Figure 5**), the hip and knee angles in the RAV display markedly broader distributions and shifts toward larger tilt values, whereas the ankle hinge remains largely unchanged. The modal hip angle increases from 12° in the isolated spike to approximately 20° in the RAV, with excursions beyond 40° (**Figure** 5). Flexibility is further amplified at the knee, whose distribution in the RAV spans 0–58° (peak at ∼28°), in contrast to the narrower range observed for the isolated spike, which rarely exceeds 35° (**Figure 5**). On the contrary, the ankle hinge remains relatively constrained, exhibiting a more prominent low-angle peak near ∼12° compared to the individual spike simulations **(Figure 5)**. This behavior indicates that the transmembrane domain and its surrounding lipids act as a mechanically conserved anchor, relatively insulated from aerosol-induced fluctuations. These results extend earlier membrane-embedded MD studies, including all-atom simulations of four glycosylated spikes in a viral membrane patch^59^, AI-driven MD of a membrane-anchored Spike–ACE2 complex ^23^, as well as single-spike membrane simulations ^58,60,61^. Here, hip and knee hinging populate larger-amplitude tilts, with the hip appearing slightly more flexible than in prior work approximating local surface crowding using a multi-spike membrane patch ^59^. This behavior is consistent with additional extrinsic interactions present in the aerosol environment (mucins, proteins, ions, and surfactant lipids), though we cannot uniquely attribute the effect to any single component. By contrast, the ankle remains a rigid pivot and shows a marked redistribution toward a low-angle mode.

**Figure 5.**
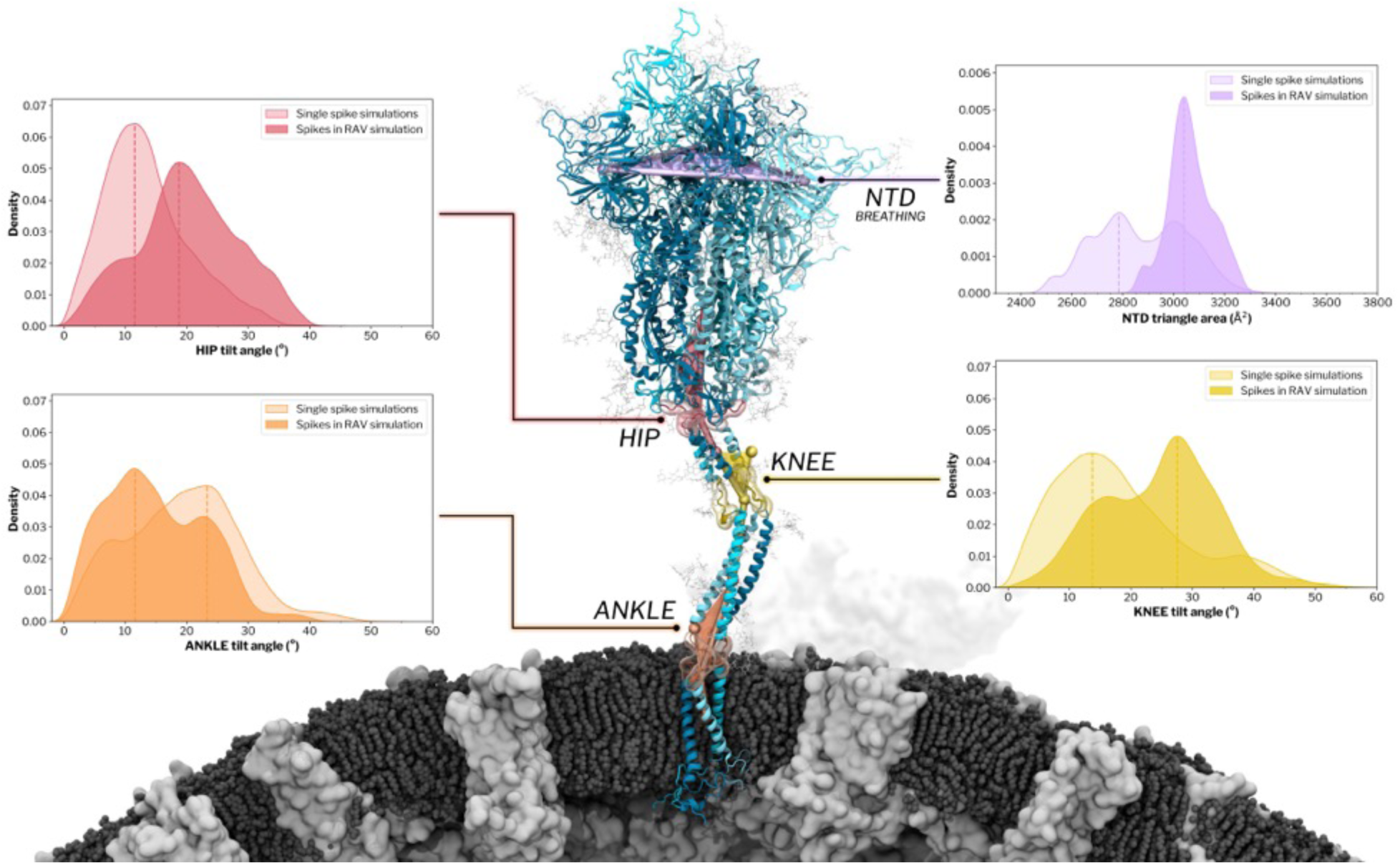
Crowded aerosol microenvironments reshape spike hinge dynamics and conformational breathing. The central panel shows a SARS-CoV-2 spike protein embedded in the intact virion from the respiratory aerosol virus (RAV) simulation, with key regions along the ectodomain and stalk highlighted: the N-terminal domain (NTD; purple), hip (red), knee (yellow), and ankle (orange). Surrounding panels report probability density distributions of the corresponding collective variables measured from MD simulations, comparing single-spike simulations from Casalino et al.^60^ (lighter shading, 10.2 μs aggregate sampling) with spikes embedded in the virion within the aerosol microenvironment (darker shading) and analyzed over the final 352 ns of production dynamics (10.2 μs aggregate sampling across 29 spikes). Left panels show hip and ankle tilt-angle distributions, while the right panels show knee tilt-angle distributions and NTD breathing quantified by the NTD triangle area. Dashed vertical lines indicate the dominant peak of each distribution.

We next assessed spike conformational breathing by tracking the triangular areas defined by corresponding domains across the three spike chains, providing a time-resolved measure of opening. The embedded Delta spike within the crowded respiratory fluid environment exhibits amplified motions in peripheral regions (S1 domain) while preserving the stability of the central core (S2 domain). The most pronounced changes occur in the N-terminal domains (NTDs), where the RAV ensemble samples NTD triangle areas approximately 150 Å^2^ larger than those observed for wild-type spikes simulated in bulk solvent, indicating an increased lateral splay of the trimer apex (**Figure 5**). Here, the NTD triangle area is defined by the planar area enclosed by connecting the centers of mass of the three NTDs and serves as a quantitative measure of spike breathing. Similar behavior has been observed in large-scale simulations of influenza virions ^24^, where hemagglutinin exhibited head “breathing” motion on a densely packed virion surface, with these motions accompanied by enhanced interplay with neighboring glycoproteins. These observations support a model in which local, interaction-mediated crowding amplifies and stabilizes breathing-like conformations through transient intermolecular contacts, consistent with the amplified NTD splay observed here in the aerosol-embedded spike ensemble. However, RBDs remain conformationally stable over the simulation timescale, retaining their initial closed or up states (20 closed and 9 up across the virion). Accordingly, the RBD–core distance distribution computed across all spikes remains bimodal and shows no significant evidence of enhanced opening/closing dynamics (**Figure S25**). We note that longer simulation timescales (on the order of hundreds of microseconds to milliseconds and outside the scope of this current work) would be required to draw conclusions about the effect of crowding on spike opening dynamics.

Overall, the enhanced flexibility at the hip and knee hinges, together with pronounced spike breathing motions, reveals a spike that becomes more dynamic in the aerosol environment while preserving the structural integrity of its fusion-competent core. Enhanced hinge flexibility and lipid shielding may influence receptor engagement probability or susceptibility to environmental stressors, suggesting a possible structural basis for experimentally observed composition-dependent infectivity differences.^13,62,63^ While such behavior contrasts with expectations derived from hard steric crowding, it is consistent with a soft, condensate-like aerosol matrix formed by multivalent assemblies, where crowding is mediated by transient, interaction-driven contacts. In such environments, coupling to a heterogeneous and dynamically fluctuating network can redistribute mechanical stress and selectively enhance conformational flexibility along compliant degrees of freedom, while leaving mechanically constrained regions unaffected ^54,55^. This framework provides a physical basis for interpreting how extreme crowding within the aerosol modulates spike mechanics: enhanced hinge-driven dynamics allow the spike to satisfy competing functional demands: maintaining an overall upright presentation during airborne transport while potentially enabling large-amplitude reorientation as the virion approaches receptor-bearing surfaces.

## 4. Conclusions

In this work, we establish an atomistic framework for understanding how the molecular organization of respiratory aerosols shapes the behavior of an airborne SARS-CoV-2 virion. Molecular dynamics simulations reveal that the aerosol rapidly organizes into a heterogeneous network of interacting components, including protein-rich assemblies, patchy surfactant lipid domains, and spatially structured ion and water distributions. This reorganization produces distinct microenvironments characterized by reduced diffusion and elevated macromolecular crowding around the virion. Mucins and albumins co-assemble into dynamic protein networks, while surfactant lipids form micelle-like clusters and surface-associated patches that collectively generate the emergent material properties of a soft, interaction-mediated aerosol matrix not captured by simplified solution-phase models.

Within this structured aerosol environment, the spike protein undergoes a selective and persistent reorganization of its local molecular neighborhood. Transient interactions with highly mobile monovalent ions are rapidly depleted, whereas divalent cations—particularly Ca²⁺—are preferentially retained at specific spike regions, including hinge sites that regulate large-scale motions. In parallel, surfactant lipids are actively recruited to the spike surface through glycan-mediated interactions, forming a heterogeneous lipid corona. This selective enrichment reflects interaction-driven partitioning within a condensate-like physical regime, rather than nonspecific accumulation driven by bulk abundance. These multicomponent interactions do not rigidify the spike; instead, they redistribute mechanical flexibility, enhancing motion at key stalk hinges while preserving a stable membrane anchor.

By framing virus-laden respiratory aerosols as chemically heterogeneous, condensate-like assemblies^53,54,56,58^, this work provides a unifying physical basis for linking aerosol composition, mesoscale organization, and viral protein dynamics. In this view, the aerosol is not a passive carrier but an active, interaction-mediated environment that conditions viral structure and mechanics during airborne transport. We show that extreme molecular crowding does not suppress viral protein motion but can instead redistribute and selectively enhance it. In addition, through resolving the molecular organization of a chemically complex respiratory aerosol, these simulations provide a structural foundation for interrogating how atmospheric chemistry and indoor air composition (e.g., trace gases and humidity) may influence viral stability and ultimately, infectivity. This molecular perspective reframes airborne disease transmission through the lens of soft-matter organization and molecular mechanics, opening new avenues for understanding—and potentially modulating—viral stability in flight.

## 5. Acknowledgements

We thank Profs. Kim Prather, Vicki Grassian, Jonathan Reid, and Linsey Marr for many rich discussions about aerosols, Prof. Ronit Freeman for discussions about mucins and cell surface glycans, and Profs. Rohit Pappu, Kresten Lindorff-Larsen, and Freeman for enlightening discussions related to condensates. We gratefully acknowledge the technical support from the NIH Resource for Macromolecular Modeling and Visualization at the University of Illinois Urbana-Champaign (NIH R24-GM145965), which provides advanced software tools and resources for molecular modeling and simulation.; this includes special thanks special thanks to John Stone and Dave Hardy for their help with NAMD software. This research was funded in part by the Air Force Office Scientific of Research (FA9550-22-1-0199 MURI 22). We also acknowledge support from the Meta-Institute for Airborne Disease in a Changing Climate at UCSD. Simulations were performed on the ORNL Summit supercomputer (project ID BIP245) and TACC Frontera supercomputer (CHE23002). CCT and MC were supported by Schmidt Sciences, LLC. REA conceived and supervised the project and acquired project resources. The construction, equilibration, and simulation of the individual components are credited as follows: CLM oversaw the E protein, NAW the M protein dimers; LC the spike protein; FLK and MAR the mucins; ACD and MAR the albumin. ACD constructed the composite virion and aerosol system. NAW refined and stabilized the composite system and performed all simulations and data management. NAW, CCT, ACD, FLK, LC, and MC performed the analysis. NAW, CCT, ACD, LC and REA wrote the manuscript. All authors were involved in editing the manuscript.

## Methods Summary

### System construction and simulation

An all-atom model of a virus-laden respiratory aerosol (RAV) was constructed by embedding a fully glycosylated SARS-CoV-2 Delta variant virion within a chemically representative respiratory fluid matrix. The virion model comprised 29 spike trimers, 360 membrane (M) protein dimers, 4 envelope (E) protein pentamers, and a lipid membrane representative of SARS-CoV-2. Spike, M, and E protein models were derived from previously validated all-atom constructs and independently equilibrated in membrane patches prior to virion assembly to ensure local relaxation of protein–lipid and protein–solvent interactions ^25,60^. The respiratory fluid matrix surrounding the virion was designed to reflect the major macromolecular and ionic components of expiratory aerosols ^33,34^.It included lung surfactant lipids (DPPC, DPPG, cholesterol), three mucin subtypes, human serum albumin, and physiological concentrations of Na⁺, K⁺, Cl⁻, Ca²⁺, and Mg²⁺ ions. Component ratios and baseline concentrations were adapted from experimental surrogate lung fluid formulations and scaled to reflect aerosol dehydration corresponding to equilibration at moderate relative humidity. Owing to packing limitations, mucins were incorporated at reduced absolute concentration while preserving their relative volume fraction and crowding behavior. The final particle composition represents one experimentally grounded realization of a virus-containing respiratory aerosol.

Virion and respiratory components were combined into a spherical particle of 270 nm diameter using a multistage packing, minimization, and equilibration protocol. Following integration of the equilibrated virion into the aerosol matrix, the system was solvated, minimized, and equilibrated to remove steric clashes and density inhomogeneities. The final RAV model comprised approximately one billion atoms. All molecular dynamics simulations were performed using the CHARMM36m force field^64,65^ and TIP3P waters^66,67^ under NVT conditions at 298 K. Production dynamics were propagated for 500 ns using a 4 fs timestep with hydrogen mass repartitioning^68^. Long-range electrostatics were treated with particle-mesh Ewald^69^, and temperature control was maintained using a Langevin thermostat. Initial equilibration steps were run on the ORNL Summit supercomputer, and the subsequent production steps were run on the TACC Frontera supercomputer. Full details of system construction, equilibration, and simulation parameters are provided in the Extended Methods (Supporting Information).

### Analysis

Trajectory analyses were designed to characterize molecular transport, self-organization, mesoscale aggregation, and spike conformational dynamics within the aerosol environment. Molecular mobility was quantified by computing self-diffusion coefficients from mean squared displacement (MSD) analyses using the MDAnalysis MSD framework^70^. For water, MSD-based diffusion coefficients were computed within spatially defined regions (voxels) of the aerosol to capture local heterogeneity in transport behavior. Diffusion analyses were also performed for ions, lipids, and proteins to relate molecular mobility to local crowding and composition. Lipid self-organization was characterized through density-based clustering and surface-coverage analyses, and viral membrane properties were examined via lipid diffusion, area-per-lipid, and membrane thickness calculations. Protein-protein organization within the aerosol was analyzed using distance-based and graph-based approaches to identify dynamic aggregation networks formed by mucins, albumins, and spike proteins. Interaction hotspots were identified from residue-level contact frequency maps computed over the MD trajectory and projected onto the spike structure. To compare local microenvironment composition using a concentration-normalized measure, we computed a preference index based on a normalized binding ratio (NBR, see Extended Methods) for each RAV component. This metric reports the relative enrichment or depletion of aerosol components within 6 Å of each protein class (29 spikes, 345 mucins, or 550 albumins). Spike conformational dynamics were quantified by measuring established hinge angles (hip, knee, and ankle) and domain-level breathing motions across all spikes over the final portion of the trajectory. All analyses were performed using in-house scripts based on VMD^71^ and MDAnalysis, which are available via a public GitHub repository (see Data and Code Availability). A full description of computational methods is provided in the Extended methods section of the Supporting Information.

