## Supporting Information for "Condensate-Like Organization in Respiratory Aerosols Modulates the Dynamics of an Airborne Virus"

***TABLE OF CONTENTS:***

***1. Extended Methods***

***1.1.* System construction and simulation**

**1.2. Analysis**

***2. Supplementary Figures***

***3. Supplementary Tables***

***4. Supplementary References***

### 1. Extended methods:

#### 1.1. System construction and simulation

Protein Models and Pre-equilibration

The SARS-CoV-2 spike ectodomain was taken from an all-atom model of the Delta variant constructed following the protocol of Casalino and co-workers, with the full glycan shield and experimentally observed mutations included in the final construct(Casalino et al., 2020). The membrane (M) protein dimer was first modelled from an ORF3a-based homology model and subsequent physics-based structure refinement procedure in collaboration with the Feig Laboratory at Michigan State University. The Delta variant of the M protein dimer was generated by introducing the I82T mutation with VMD's Mutator plugin. The envelope (E) protein pentamer was built by patching two incomplete PDB structures, 5X29(Surya et al., 2018), and 7K3G(Mandala et al., 2020), to generate the full-length model. Palmitoylation modifications were added at cysteines 40, 43, and 44, using Schrodinger's Maestro software. Each membrane protein was embedded in a lipid bilayer representative of the viral envelope and equilibrated independently prior to virion assembly. The spike, M, and E proteins were each simulated in a membrane patch for 110 ns, 700 ns, and 41 ns, respectively, allowing local relaxation of protein–lipid and protein–solvent interactions before construction of the full virion.

SARS-CoV-2 Virion Assembly

The SARS-CoV2 Delta virion (V) model was constructed following the protocol described by Casalino and co-workers(Dommer et al., 2023), using CHARMM-GUI(Jo et al., 2008), LipidWrapper(Durrant & Amaro, 2014), and Blender(Blender Online Community, 2020), a 350 Å lipid bilayer with an equilibrium area per lipid of 63 Å2 and a 100 nm diameter Blender icospherical surface mesh, The resulting lipid membrane was solvated in a 1100 Å3 waterbox and subjected to 4 rounds of equilibration and patching(Dommer et al., 2023). 360 M dimers and 4 E pentamers were then tiled onto the surface, followed by random placement of 29 full-length Spike proteins (9 open, 20 closed) according to experimentally observed Spike protein density(Ke et al., 2020). M and E proteins were oriented with intravirion C-termini. After solvation in a 1460 Å waterbox, the complete Virus model tallied >305 million atoms. The virus was equilibrated for 41 ns prior to placement in the respiratory aerosol (RA) model. The equilibrated membrane was 90 nm in diameter and remains in close structural agreement with the experimental studies(Ke et al., 2020). The final lipid membrane composition was composed of 43.0% POPC, 18.9% POPE, 6.5% POPS, 10.6% POPI, and 21.0% cholesterol.

Respiratory Aerosol Matrix Composition

The respiratory fluid matrix surrounding the virion was designed to mimic the major macromolecular and ionic components of expiratory aerosol at physiological pH. The chemical composition and component ratios for the respiratory fluid was based on the artificial saliva and surrogate deep lung fluid compositions from(Vejerano & Marr, 2018; Walker et al., 2021). The system included high–molecular weight mucins, modeled as extended, heavily O-glycosylated bottlebrush polymers, together with human serum albumin (ALB) as a representative abundant soluble protein in respiratory fluids. Albumin was taken from an experimental structure, protonated at pH 7, solvated, neutralized, minimized, and equilibrated for 7 ns prior to incorporation into the aerosol.

The total concentrations were increased by roughly 6x to estimate the environment of a particle that has reduced in diameter to roughly half of its initial size, from ~500nm to 270nm (Figure S1). The overall chemical composition and component ratios for the respiratory fluid (lipids, proteins, mucins, ions, and small solutes) were chosen based on experimental measurements of airway mucus and evaporating respiratory droplets, with modest down-sampling of the largest components (mucins) to maintain tractable system sizes while preserving their volume fraction and crowding behavior. Monovalent and divalent ions ($Na^+$, $K^+$, $Cl^-$, $Ca^{2+}$, $Mg^{2+}$) were added to reproduce near-physiological ionic strengths and charge neutrality. Final concentrations for the components in the respiratory fluid are listed in **Table S1**.

Respiratory Aerosol–Virion Assembly and Molecular Dynamics

The virion and respiratory components were combined into a single particle intended to represent a respiratory aerosol (RA) particle. All components (virion, mucins, albumin, lipids, and ions) were first placed into a 100nm cube using PACKMOL and were minimized and briefly equilibrated before being tiled into a 300nm cube. A spherical section corresponding to a diameter of 270nm was carved from this cube. An additional sphere was carved out from the inside of the particle matching the size of the SARS-CoV-2 membrane and spike proteins into which the equilibrated virus was placed. Atom clashes were resolved with a 1.2 Å cutoff. Hydrogen mass repartitioning(Hopkins et al., 2015) was applied to the structure to improve performance. The simulation box was increased to 2800 Å per side to provide a 100 Å vacuum atmospheric buffer. The system was minimized for 100K steps using conjugate gradient minimization to further reduce clashes. After minimizing, the respiratory aerosol + virus (RAV) system was heated to 298 K with 0.1 kcal/mol $Å^2$ restraints on the viral lipid headgroups, then equilibrated for 1.5 ns. Air pockets formed at the boundary of the virus integration into the aerosol due to decreased water density, which were subsequently re-solvated and equilibrated in an iterative fashion. Despite the membrane patching rounds during equilibration, a new pore formed late into the simulation and was patched at 374.6 ns. After patching the hole, the system was re-equilibrated for 2 ns and the remaining 130 ns were simulated unrestrained. These aerosol construction details were partially adapted from Dommer et al. (2023). Throughout the simulation the virus remained stable with the overall diameter of the membrane largely staying constant aside from the initial equilibration of the system and the introduction of new membrane lipids after the membrane hole was patched **(Figure S26)**. Area-per-lipids for the 5 lipid types in the membrane remain consistent across the simulation as well with all the phospholipids at around 60 $Å^2$ and cholesterol at ~40 $Å^2$ **(Figure S27)**. The aerosol's water evaporation reached an equilibrium with ~6100 water molecules in the gas phase throughout the final 120 ns of the simulation. Using the volume of the simulation box not occupied by the aerosol and the literature vapor pressure of water at 298.15 K we can get the saturated vapor density of water (Wexler & Greenspan, 1971). Comparing the observed water vapor density to this saturated vapor density we calculate an estimated equilibrated RH of 72%.

The RAV simulation was conducted in an NVT ensemble with a 4 fs timestep using the CHARMM36m force field and TIP3P water. For all simulation procedures, particle-mesh Ewald electrostatics were employed for long-range electrostatic interactions; nonbonded van der Waals interactions and short-range electrostatics were calculated with a 12 Å cutoff. The SHAKE algorithm was used to fix hydrogen bond lengths, and a Langevin thermostat with a damping coefficient of 10/ps was applied to maintain temperature control at 298.15 K. Initial equilibration steps were run on the ORNL Summit supercomputer, and the subsequent production steps were run on the TACC Frontera supercomputer utilizing 2048 computational nodes achieving a simulation speed of 7.1 ns/day. Full scaling data can be found in **Figure S28**.

#### 1.2. Analysis

<u>Lateral Diffusion of Viral Membrane Lipids</u>

To understand lipid movement in the viral membrane, we used MDAnalysis's Streamplots (Michaud-Agrawal et al., 2011) package. This generates displacement vectors across the membrane surface that represent the total motion of lipids in that location. The resulting field was given a visual representation using MAYAVI to create the 3D Streamlines representations using seed points and Blender to render the final images. The viral proteins were overlaid using the Molecular Nodes package in Blender.

<u>Identification of Viral Membrane Leaflets</u>

The inner and outer leaflets were separated by calculating the lipid's distance from the center of the virus as well as its orientation (pointing toward or away from the virion center).

Membrane Area Per Lipid (APL) Calculations

The area per lipid was calculated for the virus on a per-lipid basis by generating a spherical Voronoi tessellation with each Voronoi cell representing the given lipid's area. The center of mass of protein transmembrane regions were also included in the tessellation to best approximate the APL of the lipids surrounding membrane proteins. These points were fit to a perfect sphere with a radius given by the average radius of the points. The Voronoi representation was generated using Scipy's SphericalVoronoi package which gives the cell areas for each lipid.

Viral Membrane Thickness Measurement

Membrane thickness was measured by first separating out the inner and outer leaflets of the viral membrane and then measuring the distances between the headgroups on each leaflet. This was done on a per-lipid basis that allows us to visualize the membrane morphology in 3D.

Viral Membrane Radius Estimation

The radius of the viral membrane was calculated from the lipid head groups of either the inner or outer leaflet and the center of mass of the entire leaflet group.

Respiratory Lipid Clustering

Respiratory lipid aggregation was tracked by clustering the lipid headgroup coordinates using SciPy's DBSCAN package with the following parameters: eps = 5, min_samples = 5. The parameters were chosen based on providing qualitatively good separation between free, non-aggregated lipids and those in lipid structures. These calculations were performed every frame with a select set of timepoints visualized in **Figures 2** and **S8** along with the full timeseries in **Figure S9**.

Surface lipid clustering was tracked similarly as above with a few additional constraints. First, only lipids spatially near the surface were included in the clustering method. Second, the vectors of the lipid tail orientation were included in the clustering parameters along with the headgroup coordinates. These vectors were calculated as the vector from the COM of the head group to the COM of the tail atoms.

The surface coverage of the respiratory lipids was calculated individually for each surface cluster and then summed together. Each headgroup coordinate in the cluster was converted to polar coordinates and fit to a unit sphere after which Delauney triangulation was performed on the cluster, giving the relative surface coverage for each cluster. An example of the cluster triangulation can be found in **Figure S29** (Delauney triangulation SI). This calculation likely slightly underestimates the true surface coverage due to the method of triangulation not accounting for the actual size of the perimeter lipids and treats them as COM points. However, a rough calculation of how many lipids would be required to coat the full particle surface at an average APL of 60Å would put the final number of lipids at the surface at just under 10% of the number needed for saturation, very close the peak surface coverage calculating  which supports the fidelity of this calculation.

Spatially Resolved Water Diffusion Analysis

Diffusion coefficients for water molecules were estimated by first calculating the mean squared displacement (MSD) through the Einstein equation (Eq. S1) with MDAnalysis's MSD function. Then, on the final 10 ns of the simulation, a linear regression was fit to the "middle" region of the MSD as referenced in Maginn et. al 2019 to get the self-diffusivity coefficients as described in equation Y. For these data, the regions from 1 ns to 9 ns in the MSD were defined as the "middle" region from which the self-diffusivity coefficients were estimated. These calculations were performed on spatially segmented "voxels" within the aerosol which were cubes of 10nm x 10nm x 10nm dimensions. Regions with fewer than 1000 water molecules were discarded for having too few data points to provide accurate bulk diffusion estimates.

Eq. S1 $$MSD(r_d) = \left\langle \frac{1}{N}\sum_{i=0}^{N} \quad = |r_d - r_d(t_0)|^2 \right\rangle_{t_0}$$

Eq. S2 $$D_d = \frac{1}{(2d_\alpha)} \lim_{t\to\infty} \frac{d}{dt} MSD(r_d)$$

Pearson correlations between the calculated diffusion coefficient for a voxel and the mass of various component groups can be found in **Table S2**.

Diffusion Analysis of Aerosol and Viral Components (Ions, Lipids and Proteins)

Diffusion of all other aerosol and viral components was quantified using the same MSD-based Einstein framework described above (Water diffusion section). In-house Python scripts based on MDAnalysis were used to track particle positions along the trajectory and compute MSDs relative to reference positions at the beginning of the analyzed time window. Diffusion coefficients were obtained from linear regression of MSD versus time (Eq. S2), with the dimensionality *d* adjusted according to the nature of the motion (*d*=3 for three-dimensional diffusion and *d*=2 for lateral diffusion of membrane-embedded lipids).

**Spike protein diffusion:** Spike protein diffusion was computed independently for each of the 29 spikes present in the respiratory aerosol virus. MSDs were calculated by averaging squared displacements over all Cα atoms of a given spike. Trajectory frames were sampled at 0.5 ns intervals, and diffusion coefficients were obtained by linear regression of MSD versus time for $t \geq 20$ ns. Spike MSDs were computed from Cα atoms to capture effective mobility of the flexible ectodomain, whereas COM-based MSDs were used for other proteins to quantify translational diffusion.

**Albumin and mucin diffusion:** Diffusion of aerosol proteins was computed using center-of-mass (COM) trajectories. For each albumin molecule, the COM displacement relative to its initial position was tracked and used to compute MSDs, which were then averaged across all albumins. The same MSD framework and fitting strategy as for spike proteins was applied.

**Viral membrane protein diffusion:** Diffusion of membrane-embedded M and E proteins was quantified using COM-based MSD calculations. Although translational motion of these proteins is constrained by the viral membrane, three-dimensional MSDs were computed to enable relative comparison of protein mobility within the virion.

**Lipid diffusion:** Diffusion of respiratory (free) lipids was quantified using three-dimensional COM-based MSD analyses analogous to those used for other aerosol components. For viral membrane lipids, lateral diffusion was quantified using two-dimensional MSDs computed from lipid COM displacements projected onto the membrane surface, enabling characterization of membrane mobility and heterogeneity.

Protein aggregation and network analysis

Spatial aggregation between the three major protein classes in the SARS-CoV-2 virion and the respiratory fluid (mucins, albumin, and spike proteins) was quantified using distance-based metrics extracted from the final 500 ns of the RAV trajectory. For each frame, the center of mass (COM) of every protein segment was computed, generating a pairwise distance matrix between all selected proteins that represents the spatial organization of the system.

First, the raw distance matrix was visualized to identify large-scale proximity patterns. To resolve underlying groupings, hierarchical clustering (Ward's method) was applied to the matrix, yielding an optimized ordering of proteins that places spatially close components adjacent in the reordered matrix. The clustered distance map highlights blocks along the diagonal corresponding to persistent aggregates or semi-stable protein neighborhoods within the aerosol to identify dominant interaction patterns between protein classes.

Finally, a proximity graph was constructed to provide a complementary network-based view of aggregation. Each protein was represented as a node, and edges were drawn between protein pairs whose COM–COM distance fell below a certain cutoff. Three different cutoffs were used to describe the different aggregation layers on the RAV: 100 Å, 150 Å, and 200 Å. These cutoffs were chosen to probe aggregation at increasing mesoscale length scales. The resulting graph encodes aggregation strength as edge density, while node colors and shape denote protein class, allowing direct visualization of the mesoscale protein networks that form within the aerosol. This analysis shows the complexity of protein aggregation behavior and reveal how mucins, albumins, and spike proteins co-organize within the crowded aerosol environment.

Residue-level contact frequency analysis

To identify preferential interaction regions ("hotspots") on the SARS-CoV-2 spike protein, we performed residue-level contact frequency analyses between the spike and surrounding aerosol components along the final portion of the MD trajectory (352 ns). Contacts were identified when any heavy atom of an aerosol component was within a 6 Å cutoff distance of a spike residue. Contact counts were accumulated separately for each component type (lipids, ions, mucins, albumin) and summed over all spike protomers and all the 29 spikes present in the system.

For each spike residue, the total number of contacts was normalized by the number of analyzed frames to obtain a time-averaged contact frequency (ranging from 0, never contacted, to 1, contacted in all analyzed frames). These values report how persistently a given residue participates in interactions with the aerosol environment. Residues exhibiting consistently high contact frequencies across the trajectory were classified as interaction hotspots. Hotspot regions were visualized by mapping normalized contact frequencies onto the spike structure using a continuous color scale, allowing direct identification of spatially localized interaction patterns on the spike surface.

All hotspot analyses were performed using in-house Python scripts built on MDAnalysis and VMD-based workflows. Scripts used for contact counting and visualization are provided in the associated GitHub repository (see Data and Code Availability).

Preference Index Analysis

To quantify molecular enrichment at the spike surface while correcting for large differences in component abundance within the aerosol, we computed preference indices based on a normalized binding ratio. The calculation of the preference index follows the general logic of enrichment/depletion metrics previously used to quantify local compositional fingerprints around biomolecules, where local shell occupancy is normalized by bulk composition to correct for abundance-driven biases(Corradi et al., 2018; Shukla et al., 2009). For a given reference component (spike, mucin, or albumin), we first identified all molecules of each component type whose minimum heavy-atom distance to the reference was less than 6 Å. For each component *i*, the time-averaged number of molecules within this cutoff was computed and averaged over all copies of the reference component present in the system (e.g., 29 spikes, 550 albumins, etc).

The normalized binding ratio for component *i* was then defined as:

$$NBR_i = \frac{\langle N_i^{contact} \rangle}{N_i^{total}}$$

Where $\langle N_i^{contact} \rangle$ is the time-averaged number of molecules of component $i$ in contact with the reference, and $\langle N_i^{total} \rangle$ is the total number of that component present in the aerosol. This normalization yields a per-molecule measure of interaction propensity that is independent of absolute component abundance.

To establish a reference baseline for comparison, we computed a global mean NBR as a copy-number-weighted average over all components included in the interactome:

$$\langle NBR \rangle_{global} = \frac{\sum_i \langle N_i^{contact} \rangle}{\sum_i N_i^{total}}$$

To ensure direct comparability across reference surfaces, the global mean NBR was calculated once using all interactome datasets (spike, albumin, and mucins) and used as a fixed normalization constant for all preference index calculations.

The preference index for component i was then defined as:

$$PI_i = \frac{NBR_i}{\langle NBR \rangle_{global}}$$

Values greater than unity indicate preferential enrichment at the reference surface relative to the aerosol average, whereas values below unity indicate depletion.

This analysis was performed independently using the spike protein, mucins, and albumins as reference components, allowing direct comparison of how different biomacromolecules reshape and partition their local environments.

For visualization, we report the **preference index**, defined as the normalized binding ratio of each component divided by the global mean NBR computed across all reference systems **(Figure 4c, Figures S23-24 and Tables S3-S6)**. Bar and circular plots therefore show **relative enrichment/depletion (PI)** rather than raw contact counts or raw NBR values.

Spike Protein Tilting, Conformational Breathing, RBD openness

Spike's hip, knee, and ankle hinge angles were calculated using in-house Tcl scripts executed within VMD(Humphrey et al., 1996). The hip angle is defined by three points corresponding to (i) the center of mass (COM) of residues 816–1135 in the spike head region, (ii – vertex1) the COM of residues 1136–1140 at the head–stalk junction, and (iii – vertex2) the COM and principal axis direction of the upper stalk segment (residues 1141–1161). The knee angle is defined by three points corresponding to (i) the COM and principal axis direction of the upper stalk (residues 1141–1161), (ii – vertex1) the COM of residues 1161–1168 at the junction between the two stalk legs, and (iii – vertex2) the COM and principal axis direction of the lower stalk segment (residues 1169–1206). The ankle angle is defined by three points corresponding to (i) the COM and principal axis direction of the lower stalk (residues 1169–1206), (ii – vertex1) residue 1213 at the membrane-proximal junction, and (iii – vertex2) the COM and principal axis direction of the membrane-proximal stalk segment (residues 1212–1239).

Spike conformational breathing was quantified by computing the triangular area defined by the N-terminal domains (NTDs) across the three protomers of each spike trimer. The NTD triangle area is defined by three points corresponding to the centers of mass (COMs) of the NTDs from each spike chain, calculated over Cα atoms of residues 13–291 for chains A, B, and C, respectively. Specifically, COMs were computed for residues 13–291 in chain A, chain B, and chain C of each spike, and the planar area enclosed by connecting these three COMs was evaluated at each trajectory frame. This geometric measure provides a time-resolved descriptor of lateral opening ("breathing") of the trimer apex.

Both hinge-angles and NTD triangle area distributions were computed across all 29 spikes embedded in the quasi-spherical virion over the final 1760 frames (~352 ns) for each spike to enable direct comparison with previously reported single-spike simulations of both closed and one-RBD-up conformations(Casalino et al., 2020). This analysis window corresponds to an aggregate sampling time of approximately ~10.2 μs across all spikes.

RBD openness was assessed by measuring the distance between each receptor-binding domain (RBD) and the spike core, which reports the radial displacement of the RBD relative to the trimer core and serves as a proxy for RBD opening (Sztain et al., 2021). The RBD–core distance is defined by two points corresponding to (i) the center of mass (COM) of the RBD β-sheet region and (ii) the COM of the central helical core of the spike. RBD COMs were computed over Cα atoms of residues 375–380, 394–404, 431–438, and 508–517 for each of the three protomers (chains A, B, and C), whereas the spike core COM was computed over Cα atoms of residues 747–784, 946–967, and 986–1034. To assess temporal convergence and potential drift, distance distributions for each RBD chain were additionally evaluated over four equal trajectory segments of 632 frames each (approximately 16 ns per window).

### 2. Supplementary Figures

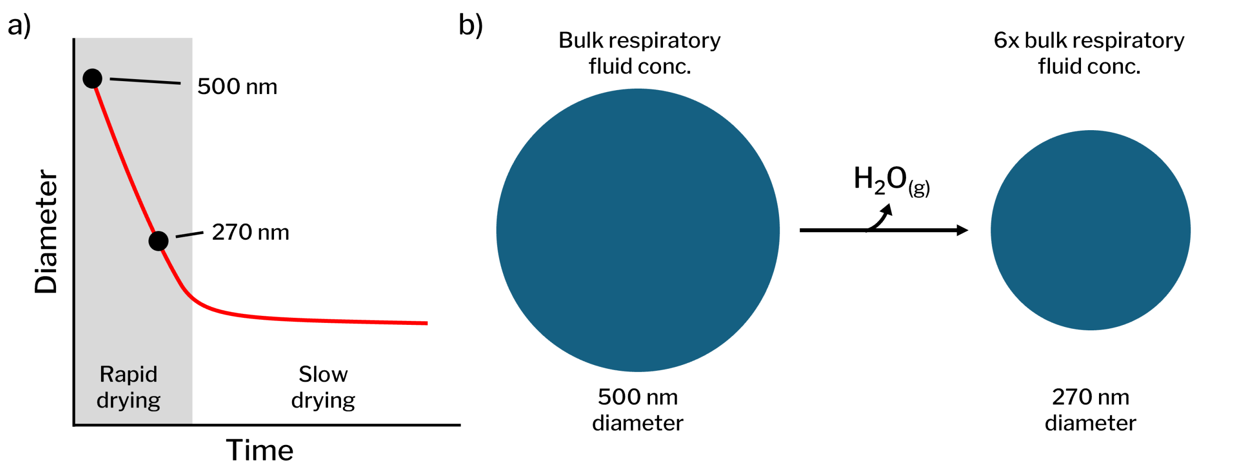


**Figure S1. Aerosol evaporation scheme.** (a) Representative plot showing the expected changes in aerosol size after emission based on the relative solute concentration increase of the simulated model. (b) Diagram of the initial emitted aerosol size and the final aerosol model with a ~6x solute concentration increase relative to bulk respiratory fluid.

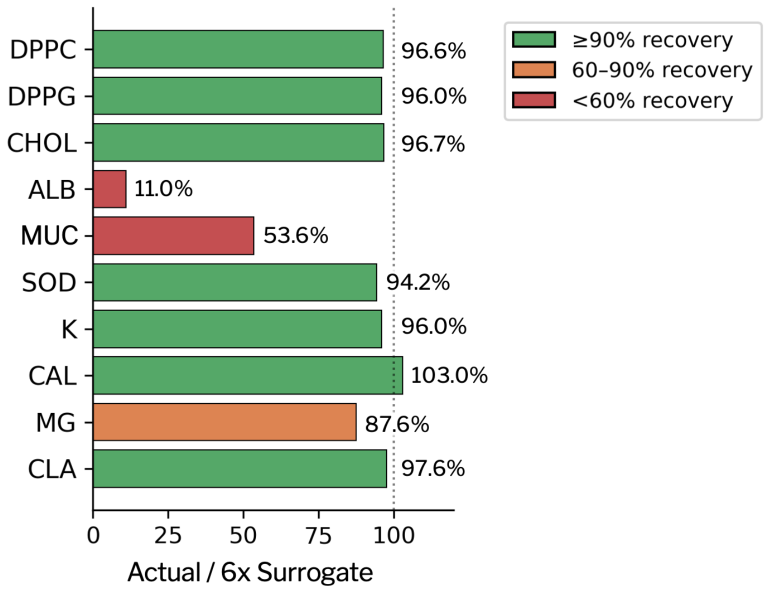


**Figure S2. Recovery of aerosol components relative to intended surrogate concentrations.** Comparison between the final RAV composition ("actual") and the target concentrations corresponding to 6× concentration of the lung fluid surrogate. Bars show percent recovery for each component relative to (left) the intended 6× concentration. Components are color-coded as ≥90% recovery (green), 60–90% recovery (orange), and <60% recovery (red). Simulation component labels correspond to: ALB (human serum albumin), MUC (respiratory mucins), DPPC (dipalmitoylphosphatidylcholine), DPPG (dipalmitoylphosphatidylglycerol), CHOL (cholesterol), SOD ($Na^+$), CAL ($Ca^{2+}$), MG ($Mg^{2+}$), K ($K^+$), and CLA ($Cl^-$). Most lipid and ionic species approach ≥90% recovery relative to the target.

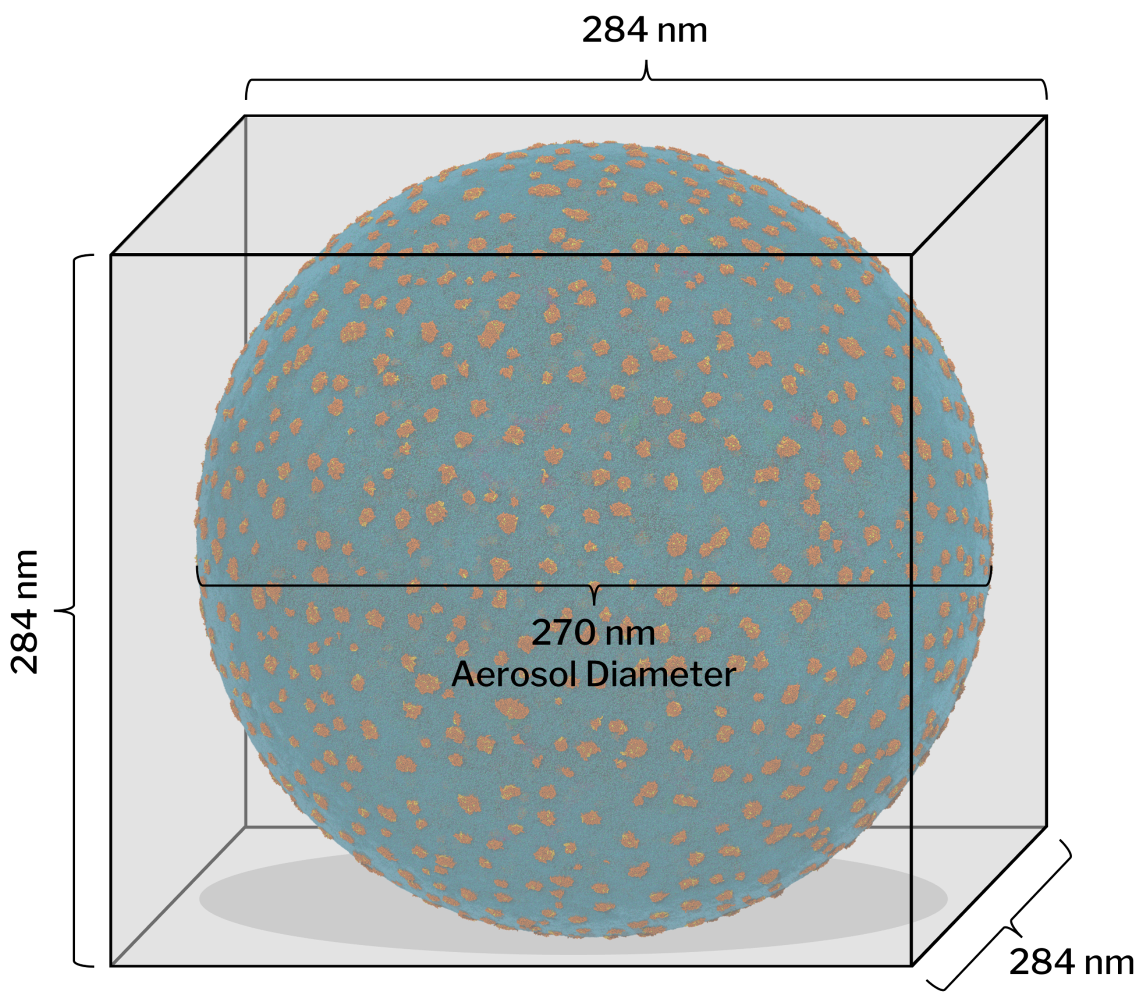


**Figure S3: Periodic boundary conditions for the aerosol system.** The aerosol has at minimum a 10 nm buffer distance to the edge of the simulation box. The region surrounding the aerosol was initialized with vacuum and as the simulation progressed, water from the aerosol formed an equilibrium with the gas phase.

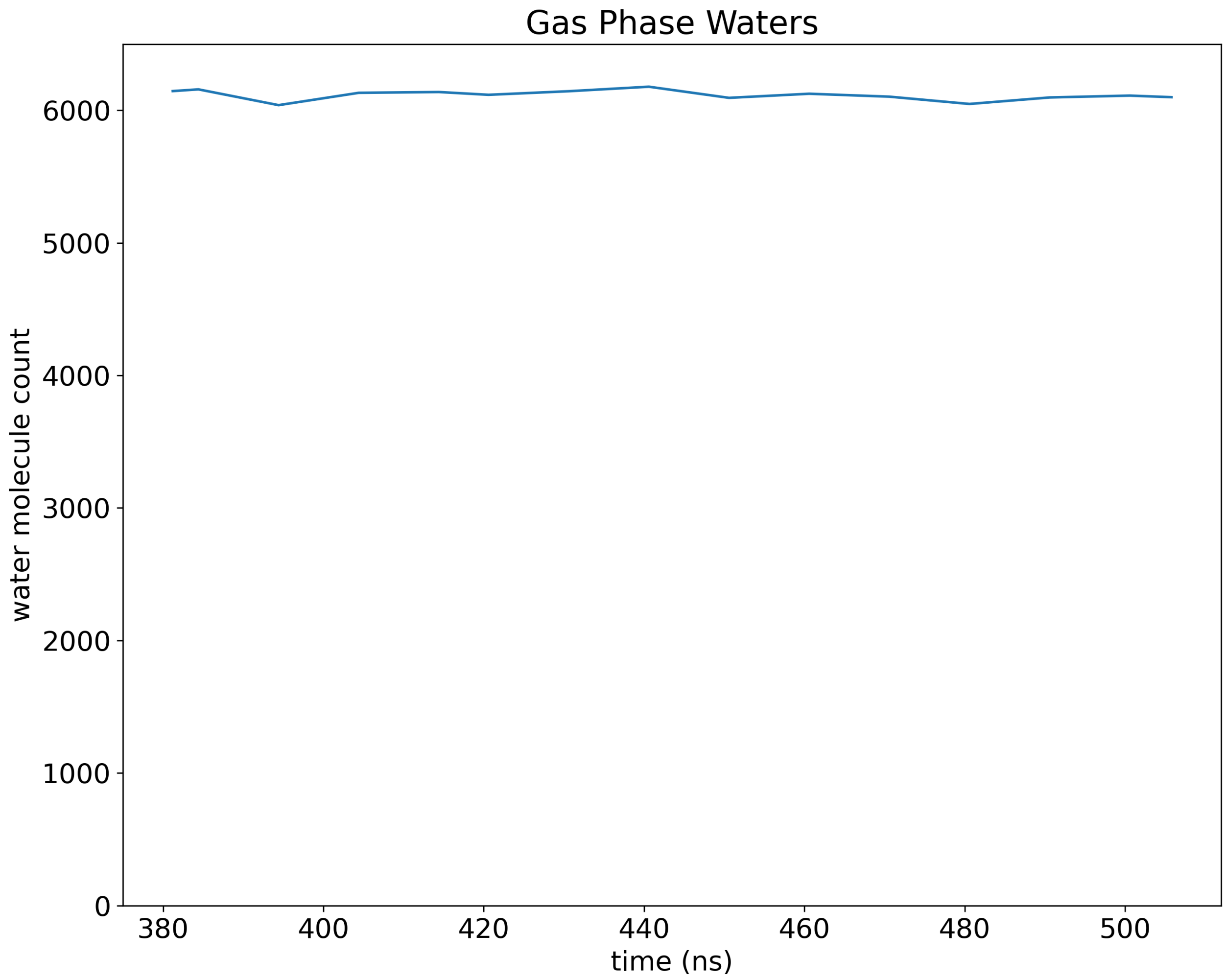


**Figure S4. Gas-phase water population during equilibration of the respiratory aerosol.** Number of water molecules located in the vacuum (gas-phase) region over the final 120 ns of the 500 ns production simulation. The gas-phase water population stabilizes at ~6,100–6,200 molecules, with only minor fluctuations over time. The absence of systematic drift indicates that a dynamic equilibrium between evaporation and condensation has been established across the gas–liquid interface.

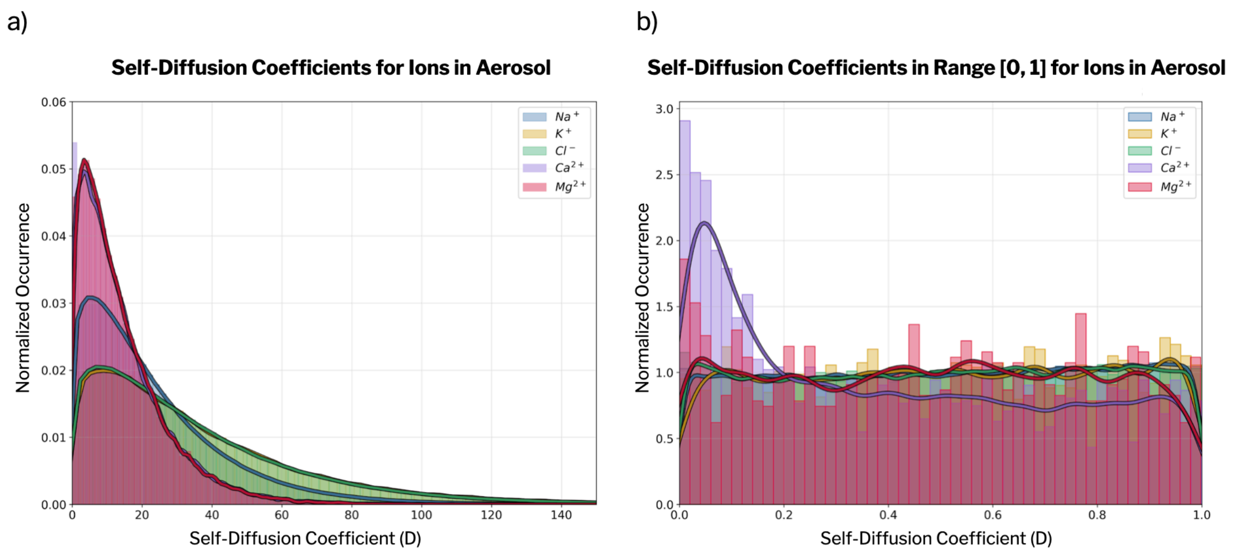


**Figure S5. Distribution of self-diffusion coefficients for ions in the respiratory aerosol.** Normalized histograms of self-diffusion coefficients (D) for the ions present in the system ($Na^+$, $K^+$, $Cl^-$, $Ca^{2+}$, and $Mg^{2+}$) computed from the 500 ns RAV trajectory using mean-squared displacement analysis (Methods). The left panel (a) shows the full distribution of diffusion coefficients, highlighting broad heterogeneity and long tails associated with highly mobile ions. The right panel (b) presents a close-up of the low-diffusion regime (range [0, 1]), enabling direct comparison of the relative populations of slow-diffusing ions. Curves correspond to kernel density estimates over the normalized histograms. The self-diffusion coefficients for ions reveal differences between monovalent and divalent ions. Monovalent ions such as $Na^+$, $K^+$, and $Cl^-$ exhibit higher diffusion coefficients, aligning with trends reported for these ions in aqueous solutions. Divalent ions, including $Ca^{2+}$ and $Mg^{2+}$, display lower diffusion coefficients, particularly for calcium. This deviation suggests persistent interactions between $Ca^{2+}$ and other components, potentially driven by its higher charge density and propensity to form longer-lasting interactions within the aerosol environment components.

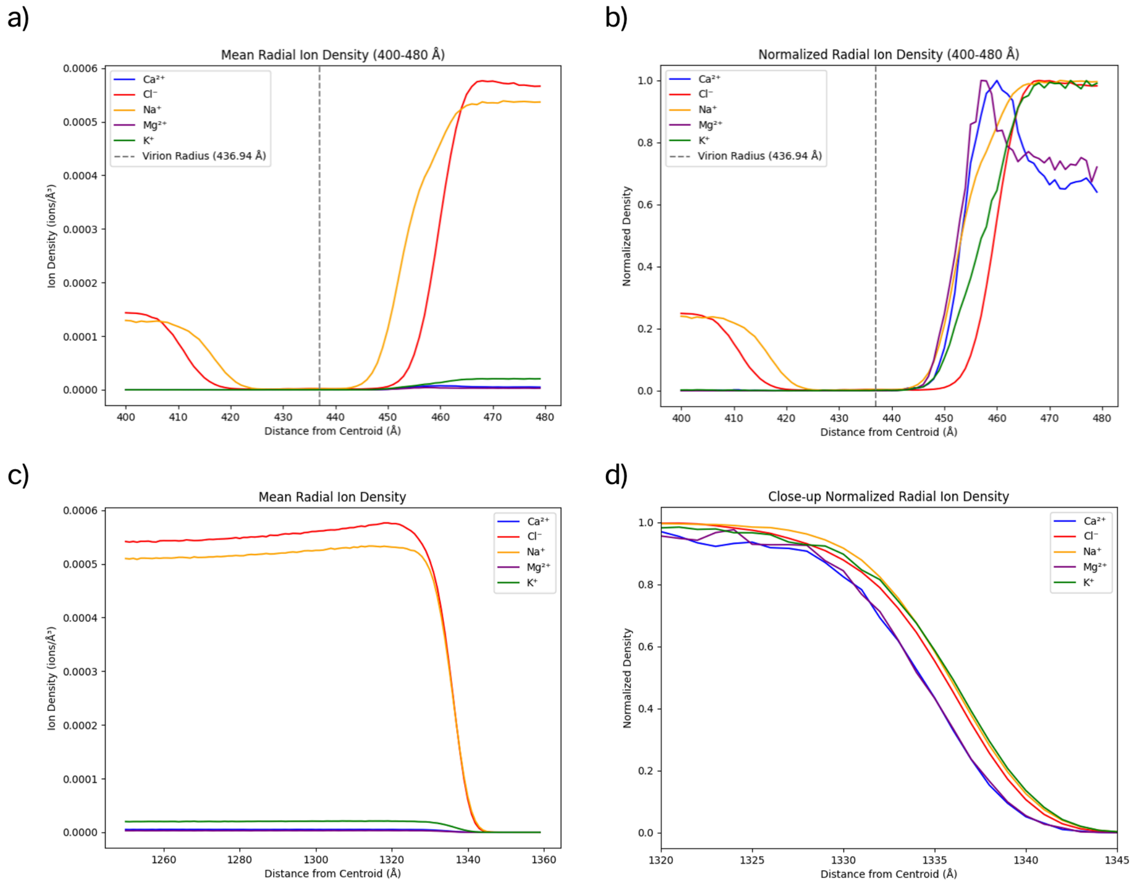


**Figure S6. Radial organization of ions around the virion and within the respiratory aerosol.** (a) Mean Radial ion density distribution and (b) Normalized mean radial ion density distribution for $Na^{+}$, $K^{+}$, $Cl^{-}$, $Ca^{2+}$, and $Mg^{2+}$ computed relative to the virion centroid and averaged over frames from 250 to 500 ns (sampled every 10 frames). Profiles are shown for the 400–480 Å range encompassing the virion surface; the dashed line indicates the mean virion radius (436.94 Å). Divalent cations ($Ca^{2+}$ and $Mg^{2+}$) are enriched in the vicinity of the virion membrane, while monovalent ions display broader distributions extending toward larger radial distances. (c) Normalized mean radial ion density distributions computed relative to the aerosol centroid, averaged over frames from 250 to 500 ns (sampled every 10 frames), shown for the 1250–1360 Å range encompassing the aerosol–air interface. Monovalent ions ($Na^{+}$, $K^{+}$, $Cl^{-}$) preferentially populate more external radial positions compared to divalent cations. (d) Close-up view of the aerosol–air interfacial region highlights a stratified ionic organization, with monovalent cations occupying the outermost layers and divalent cations enriched deeper within the aerosol, consistent with charge, hydration, and ion–lipid interaction effects.

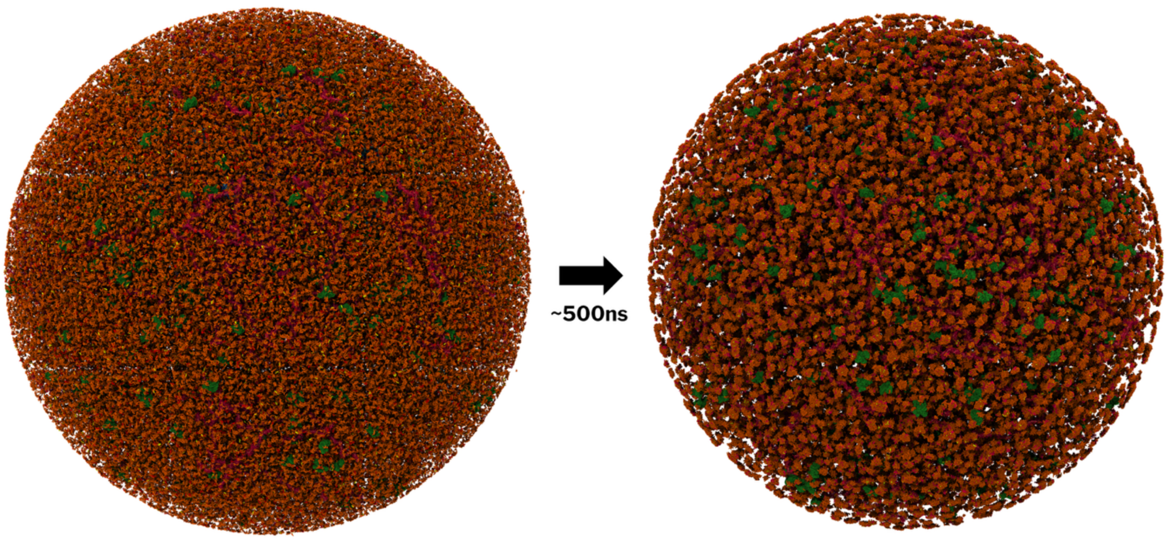


**Figure S7. Respiratory Lipid Organization.** Renders highlighting the difference in lipid structure from the start of the simulation (left) to the last frame (right). (Orange:Lipids, Green:Albumin, Red:Mucin)

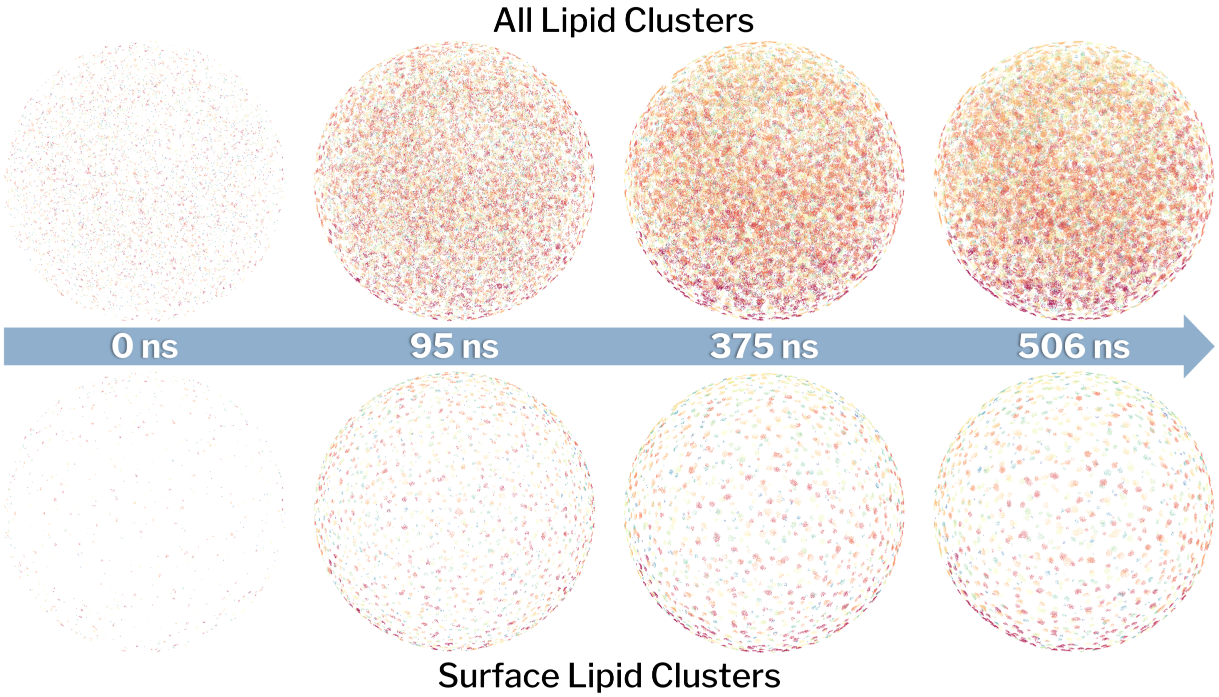


**Figure S8. Plots highlighting the change in respiratory lipid clusters across the simulation.** These plots represent the total lipid clusters for the full system (top) and just the surface (bottom) at different timepoints in the simulation. Each of the clusters is colored differently by cluster number to help differentiate between clusters.

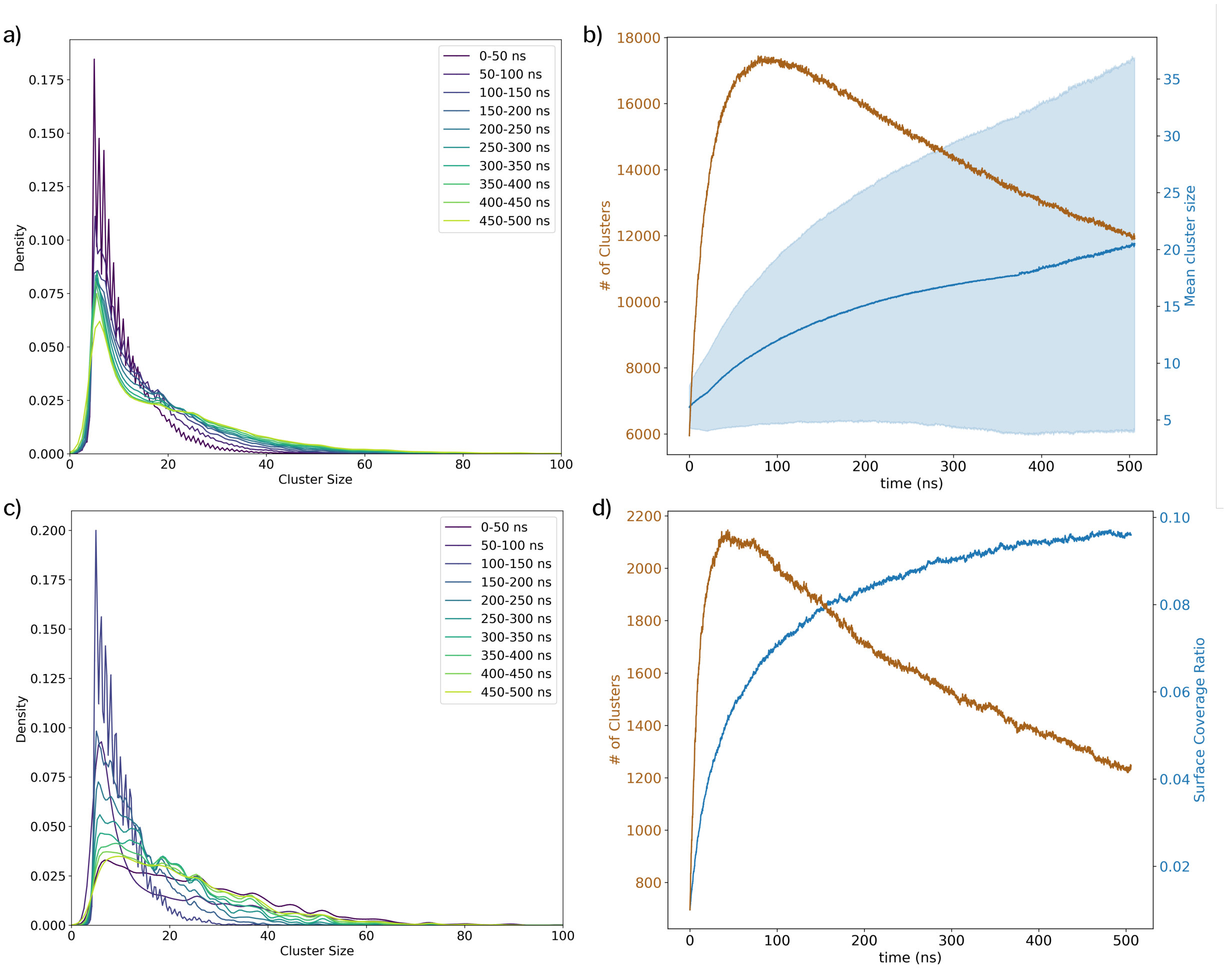


**Figure S9. Time-resolved lipid clustering dynamics in the respiratory aerosol virus (RAV).** (a) Probability density distributions of lipid cluster sizes computed over successive 50 ns windows across the 500 ns production trajectory for the full system. Over time, the distribution progressively shifts toward larger cluster sizes, indicating coarsening of lipid assemblies. (b) Time evolution of the total number of lipid clusters (brown, left axis) and the mean cluster size (blue, right axis). An initial rapid increase in cluster number is followed by a gradual decrease, while the mean cluster size steadily grows, consistent with cluster merging and reorganization. The blue shaded region represents ±1 standard deviation of the mean cluster size.(c) Cluster size distributions for the same trajectory windows emphasizing the persistence and growth of intermediate-to-large lipid aggregates at later times. (d) Time evolution of the total number of clusters (brown, left axis) and the lipid surface coverage ratio (blue, right axis). While the number of clusters decreases after the initial nucleation phase, the surface coverage increases monotonically, reflecting the formation of larger, more contiguous lipid domains.

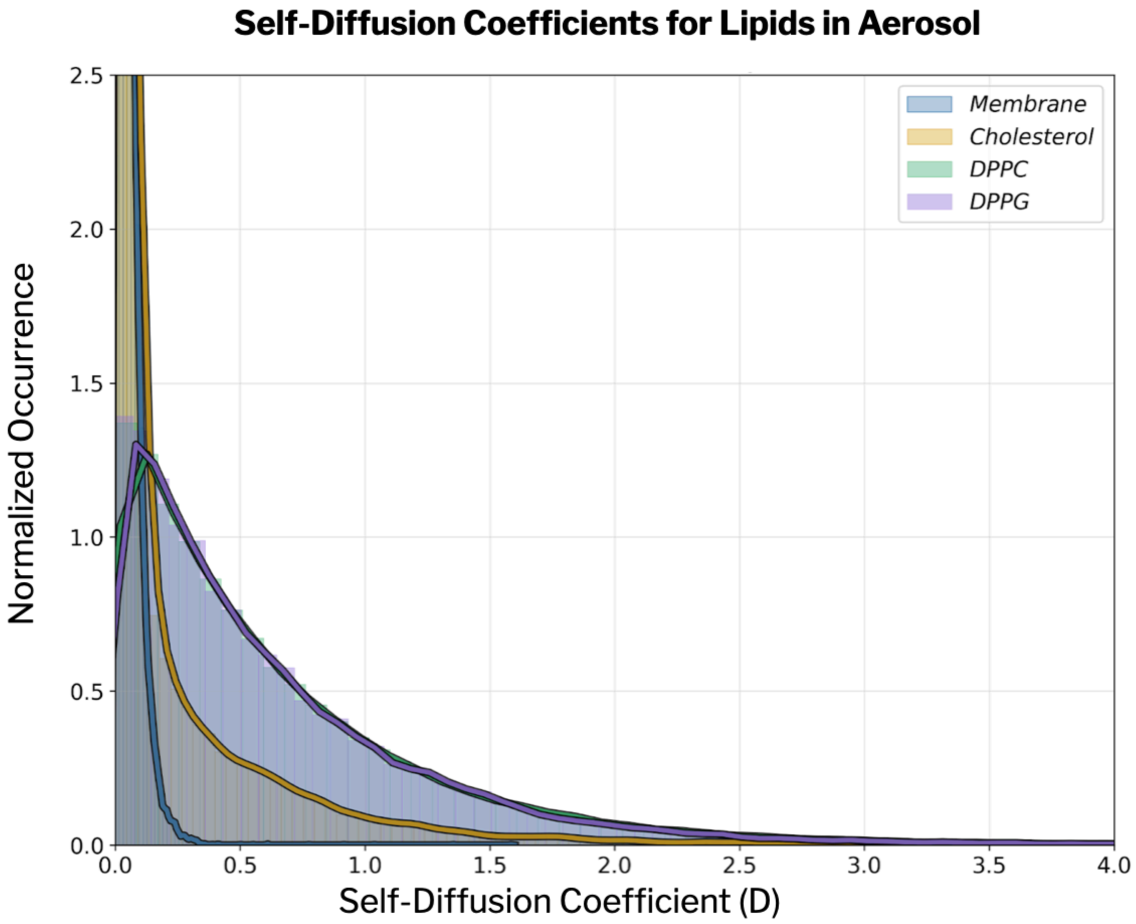


**Figure S10. Distribution of self-diffusion coefficients for lipids in the respiratory aerosol.** Normalized histograms of self-diffusion coefficients (D) are shown for viral membrane lipids and aerosol-associated lipids (DPPC, DPPG, and cholesterol). Membrane lipids exhibit low diffusion coefficients, reflecting their structural role in maintaining membrane integrity and the constrained environment of the viral envelope. In contrast, aerosol lipids display markedly higher diffusion coefficients and broader distributions, indicative of increased mobility within the aerosol phase. Differences among aerosol lipid species further reflect their distinct chemical properties and interactions with surrounding proteins, ions, and water.

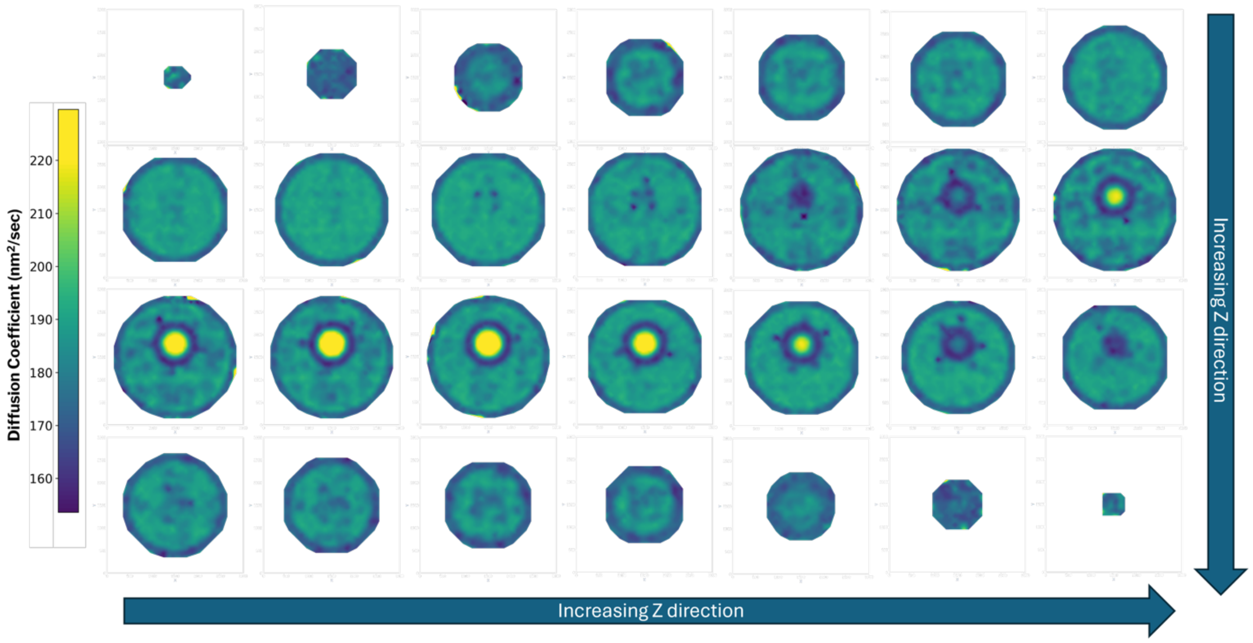


**Figure S11. Spatial heterogeneity of water diffusion across the respiratory aerosol.** Two-dimensional contour maps of local water diffusion coefficients computed using voxel-based mean-square displacement analysis are shown for consecutive 10 nm–thick slices along the Z direction. Diffusion values are color-coded, with higher mobility shown in yellow and lower mobility in blue. The bright central region corresponds to the virion interior, which contains only water and NaCl and therefore serves as an internal bulk-like reference for water diffusion. Reduced diffusion is observed near the air–water interface and in crowded regions surrounding the virion. A three-dimensional rendering of the full diffusion volume is provided in **Movie M1**.

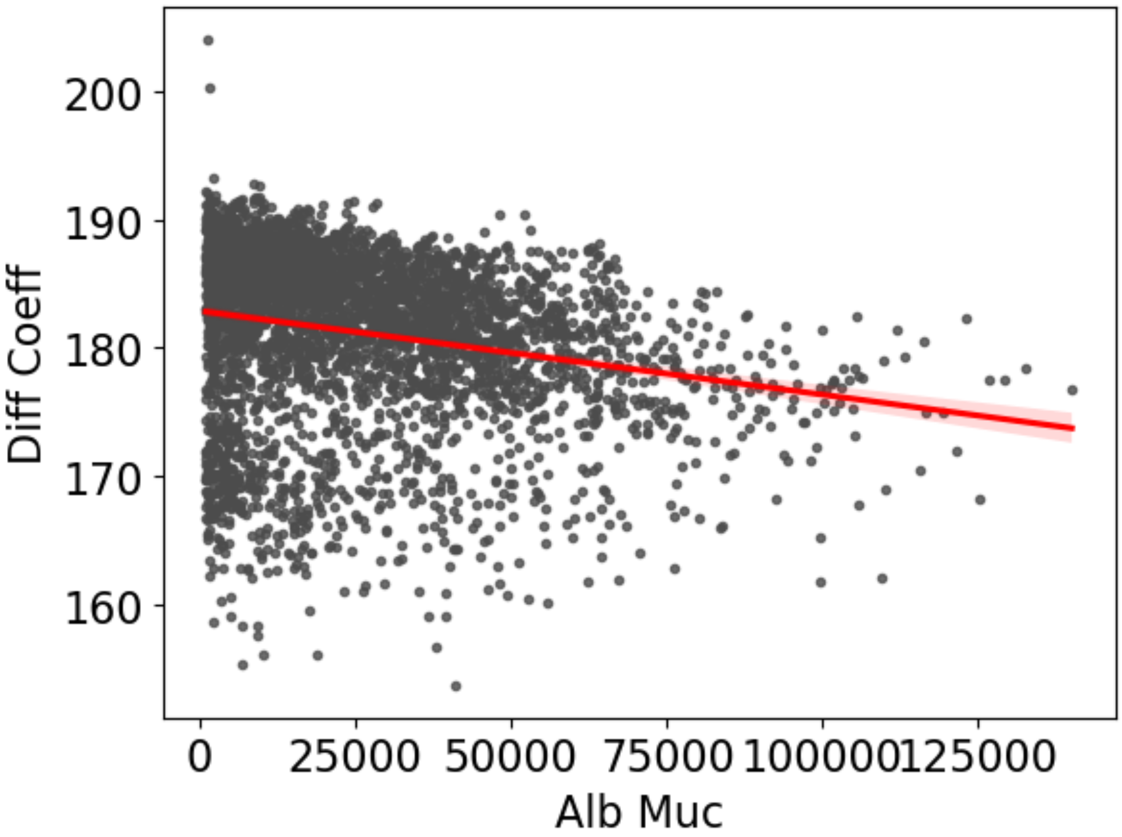


**Figure S12. Relationship between local water diffusion and protein crowding in the respiratory aerosol.** Local water diffusion coefficients are plotted as a function of increasing combined mucin and albumin content within aerosol voxels. Each point represents a spatial voxel averaged over the analyzed trajectory window. A first-order linear regression is shown in red, with the shaded region indicating the 99% confidence interval. The trend indicates reduced water mobility in regions with higher protein density, consistent with transport restrictions imposed by macromolecular crowding.

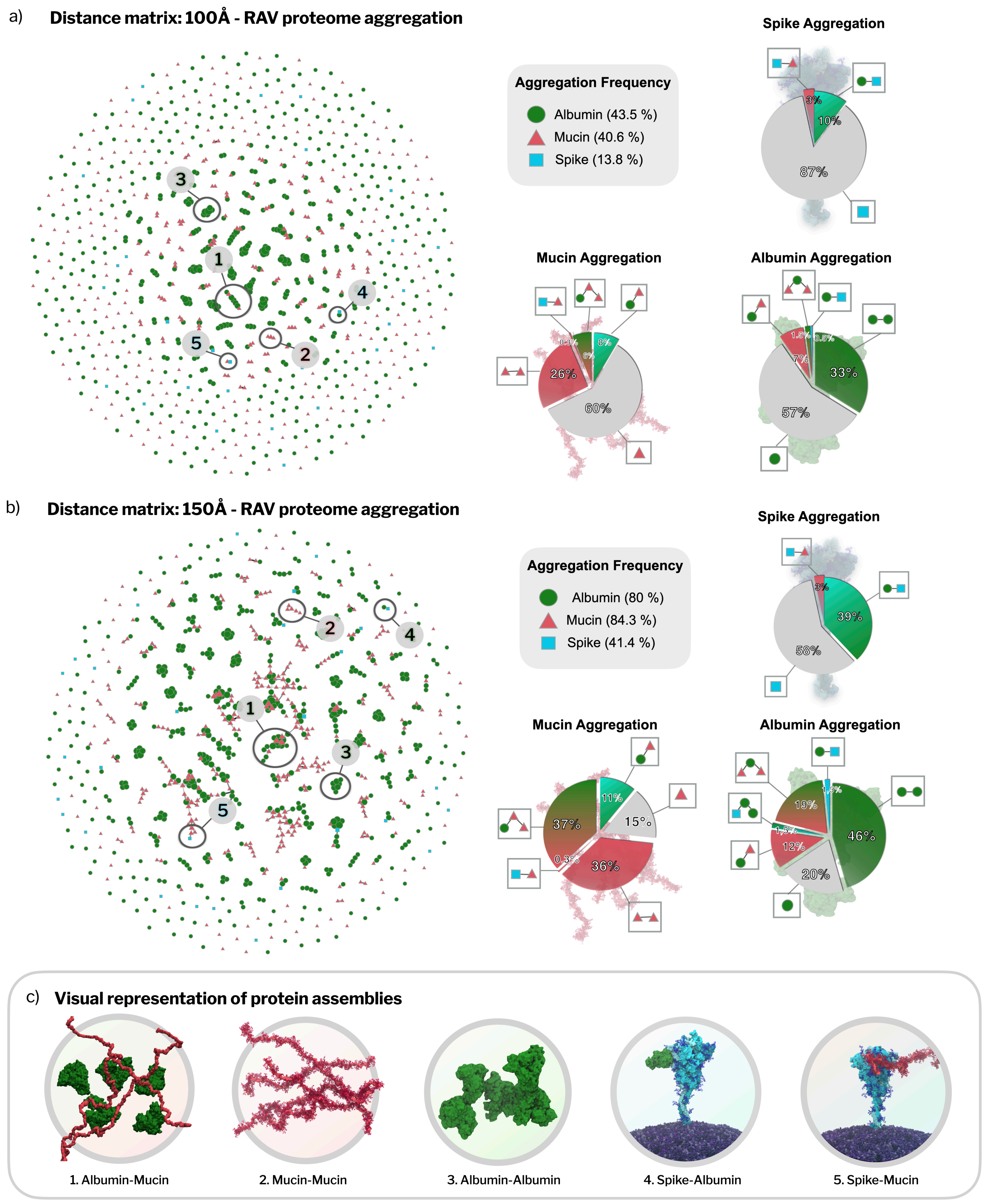


**Figure S13. Distance-threshold sensitivity of RAV proteome aggregation.** Distance-dependent aggregation of aerosol and virion proteins evaluated at progressively permissive COM distance cutoffs. (a) RAV proteome aggregation at a 100 Å cutoff. Nodes represent individual proteins (albumin, mucin chains, and spike trimers), and edges connect protein pairs whose centers of mass lie within 100 Å. Connected components define "assemblies." The aggregation frequency panel reports the fraction of each protein type participating in assemblies under this strict distance criterion. Pie charts summarize the composition of assemblies centered on each protein type (spike, mucin, and albumin). (b) RAV proteome aggregation at a 150 Å cutoff. Increasing the cutoff results in larger and more interconnected assemblies, reflected by increased aggregation frequencies and shifts in assembly composition toward mixed protein clusters. (c) Representative structural examples of protein assemblies identified under the distance-based criterion, illustrating albumin–mucin, mucin–mucin, albumin–albumin, spike–albumin, and spike–mucin associations. Together, these analyses demonstrate that aggregate size and compositional heterogeneity increase with distance threshold, while preserving the dominant trends in mucin- and albumin-driven network formation.

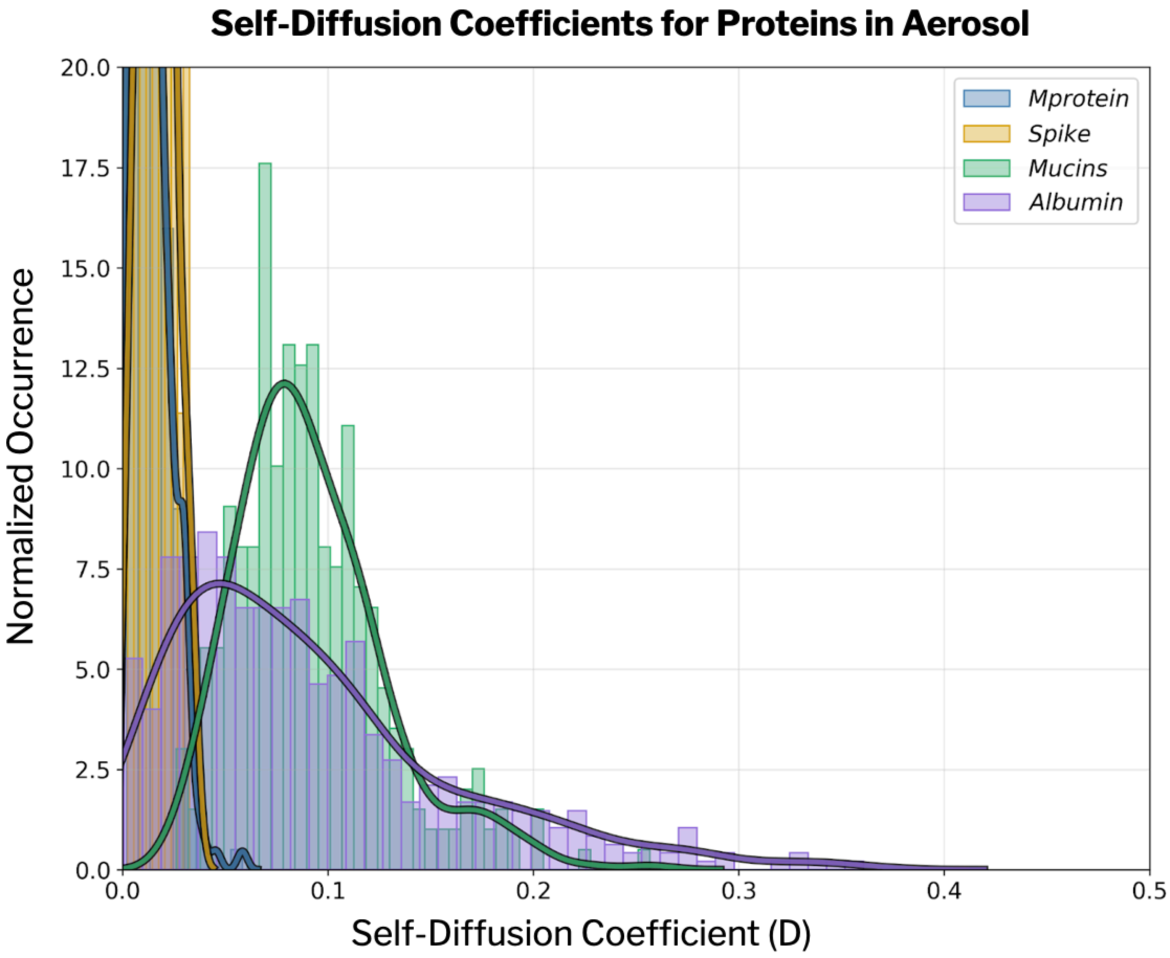


**Figure S14. Distribution of self-diffusion coefficients for proteins in the respiratory aerosol.** Normalized histograms of self-diffusion coefficients (D) are shown for membrane-associated proteins (spike protein and M protein) and aerosol proteins (mucins and albumin), computed from the 500 ns RAV trajectory using mean-squared displacement analysis. Membrane-bound proteins exhibit strongly restricted diffusion, consistent with anchoring to the viral membrane and limited translational mobility. In contrast, aerosol proteins display substantially broader diffusion distributions. Mucins show a wide range of diffusion coefficients, reflecting their heterogeneous conformations and participation in extended, transient networks. Albumin exhibits overall higher mobility than membrane-bound proteins, while still displaying a low-diffusion tail indicative of frequent interactions with other aerosol components.

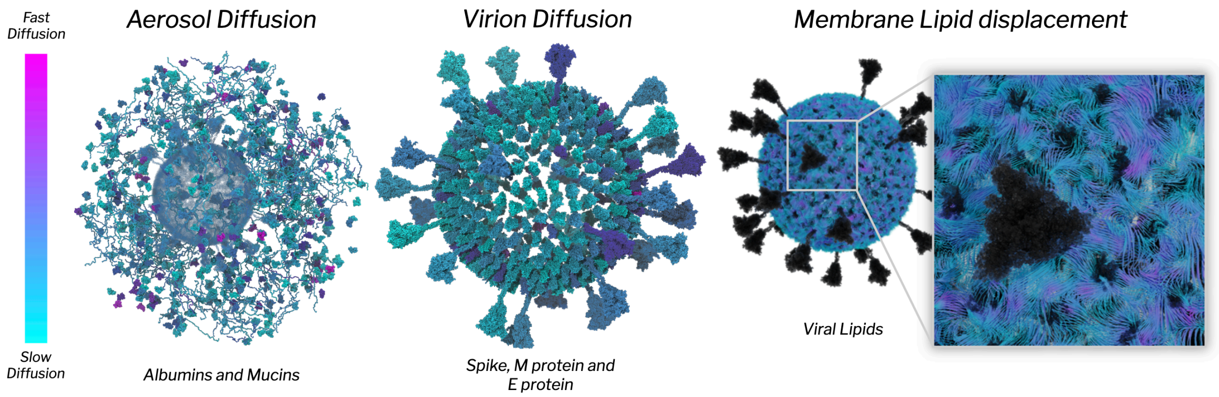


**Figure S15. RAV component diffusion.** Self-diffusion coefficients colored mapped onto representative macromolecular components in the aerosol (albumin and mucins; left) and virion membrane-embedded proteins (spike, M, and E; middle). Right: membrane lipid displacement in the viral envelope, with a zoomed region illustrating local rearrangements within the membrane. Viridis scale denotes relative diffusion (slow (green) to fast (purple)).

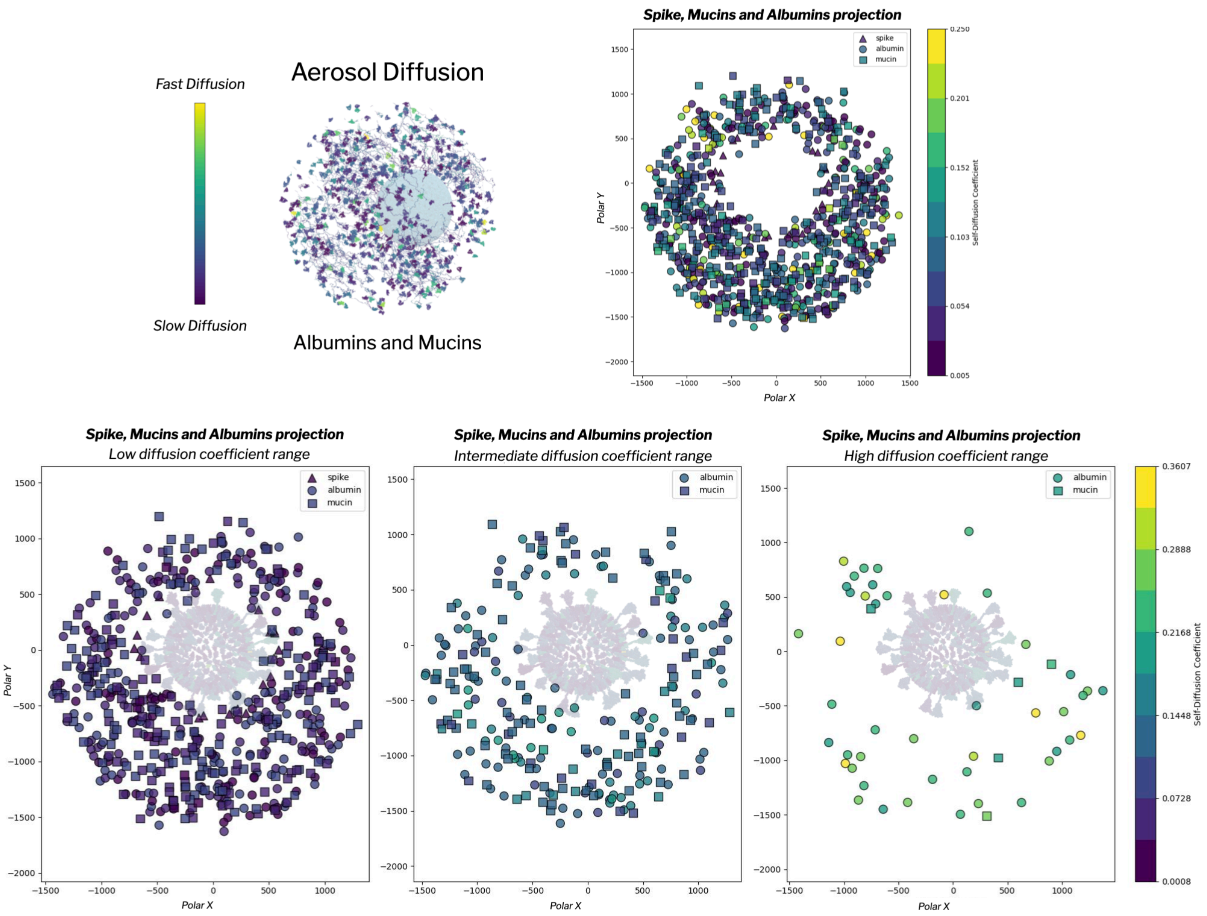


**Figure S16. Spatial organization of protein self-diffusion within the respiratory aerosol.** Protein self-diffusion coefficients are mapped onto spatial projections of the respiratory aerosol to visualize how local environment influences macromolecular mobility. Proteins are shown in a polar XY projection, with radial distance corresponding to the three-dimensional distance from the aerosol center and colors indicating self-diffusion coefficients (purple, slow diffusion; yellow/green, fast diffusion). Proteins located toward the aerosol periphery exhibit higher diffusion coefficients, consistent with reduced crowding and increased exposure to water-rich or interfacial regions. In contrast, proteins with lower diffusion coefficients cluster preferentially toward the aerosol interior and around the central viral structure, where macromolecular crowding and frequent protein–protein or protein–lipid interactions restrict translational motion. Subsets of proteins participating in larger or denser aggregates display systematically reduced diffusion, highlighting the role of transient network formation in shaping spatially heterogeneous transport within the aerosol.

**Macromolecular and Membrane Diffusion in the RAV.** To complement the voxel-based transport analysis of water and small solutes, we examined the mobility of macromolecular components within the respiratory aerosol virus (RAV). Self-diffusion coefficients were computed for representative aerosol proteins (albumin and mucins), virion membrane proteins (spike, M, and E), and viral membrane lipids, and mapped onto structural representations of the system (**Figure S15**).

Aerosol proteins exhibit spatial mobility gradients similar to those observed for water. Proteins located near the aerosol periphery display relatively higher diffusion coefficients, whereas those embedded in interior, protein-rich regions exhibit reduced mobility. This trend reflects increased macromolecular crowding and intermolecular contacts toward the particle core, which constrain translational motion. Within the virion, membrane-associated spike and M proteins exhibit substantially lower diffusion compared to free aerosol proteins, consistent with their anchoring in the viral envelope. Viral membrane lipids also diffuse more slowly than free respiratory lipids in the aerosol phase, reflecting confinement to two-dimensional lateral motion and bilayer packing constraints. To further characterize membrane dynamics, lipid trajectories were used to generate displacement streamlines over the course of the simulation (**Figure S15, right**). These reveal spatial heterogeneity in lateral lipid mobility, with localized regions of enhanced diffusion. Faster lipid motion frequently occurs adjacent to membrane proteins and along their contours, indicating that local protein–lipid interactions modulate membrane fluidity. The spatial scale and transient nature of these features are inconsistent with large-scale raft-like domain formation and instead suggest dynamic, protein-driven lipid reorganization at the virion–aerosol interface.

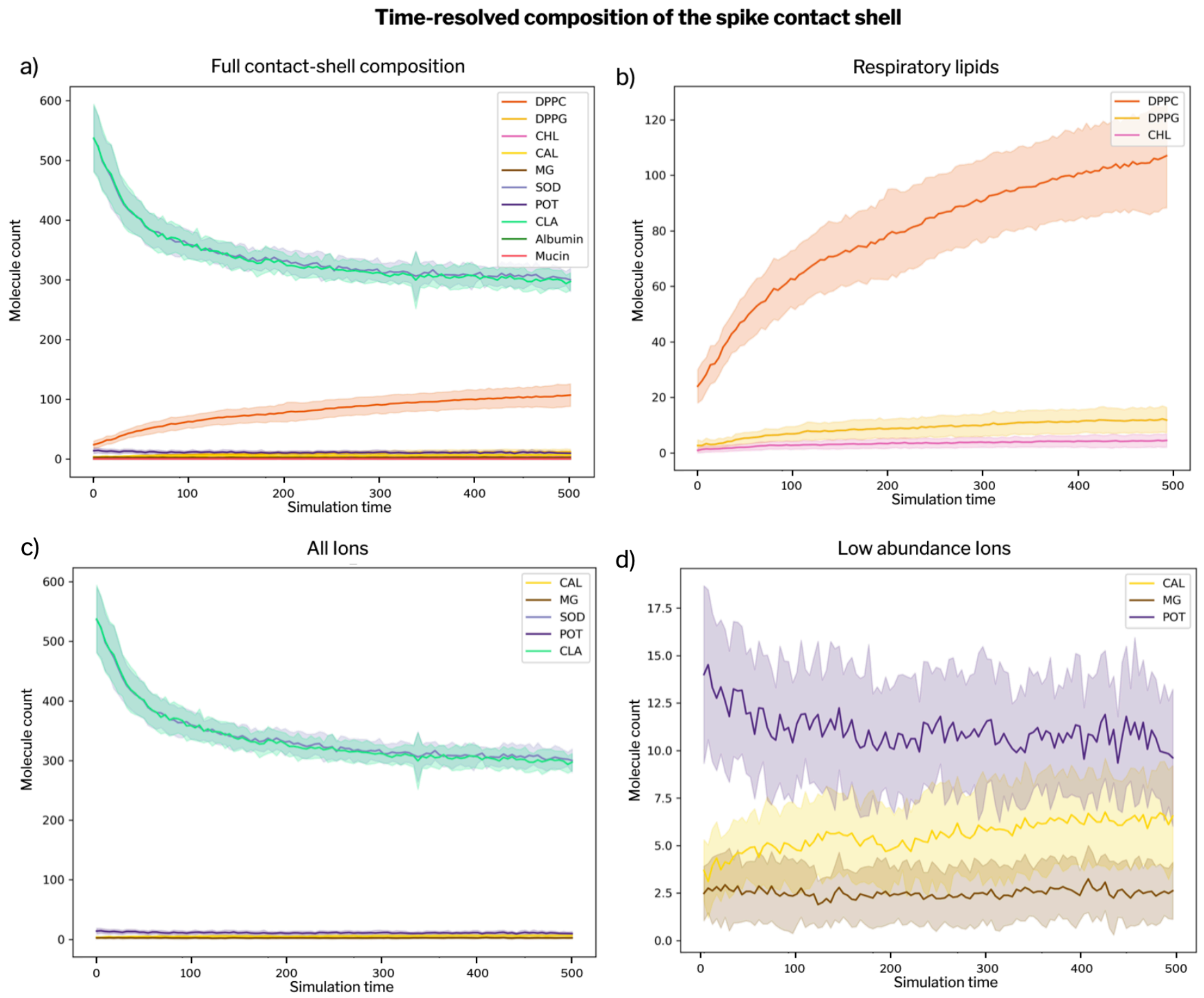


**Figure S17. Time-resolved composition of the spike contact shell in the RAV.** Mean (solid lines) and standard deviation (shaded regions) of the number of molecules within the 6 Å spike contact shell, averaged over all spikes, as a function of simulation time. a) Full contact-shell composition. Time evolution of all quantified components, including respiratory lipids (DPPC, DPPG, CHL), proteins (Albumin, Mucin), and ionic species (SOD, CLA, CAL, MG, POT). A pronounced early depletion of monovalent ions ($Na^+$ and $Cl^-$) is observed, followed by stabilization, while respiratory lipids progressively accumulate within the spike contact shell. b) Respiratory lipids. Detailed view of DPPC, DPPG, and CHL dynamics. DPPC exhibits a sustained and monotonic increase, whereas DPPG and cholesterol show more modest enrichment over time. c) All ions. Evolution of ionic species in the spike microenvironment. $Na^+$ (SOD) and $Cl^-$ (CLA) display rapid early depletion and subsequent equilibration, whereas divalent cations and $K^+$ remain comparatively stable. d) Low-abundance ions. Zoomed view of $Ca^{2+}$ (CAL), $Mg^{2+}$ (MG), and $K^+$ (POT) dynamics. $Ca^{2+}$ shows a mild increase over time, while $Mg^{2+}$ and $K^+$ fluctuate around approximately constant mean values, indicating limited long-timescale redistribution. Component names correspond to their identifiers in the simulation system: SOD = $Na^+$, CLA = $Cl^-$, CAL = $Ca^{2+}$, MG = $Mg^{2+}$, POT = $K^+$, and CHL = cholesterol. Together, these data reveal an initial ion reorganization phase followed by progressive lipid enrichment, highlighting the dynamic restructuring of the spike's local microenvironment during aerosol equilibration.

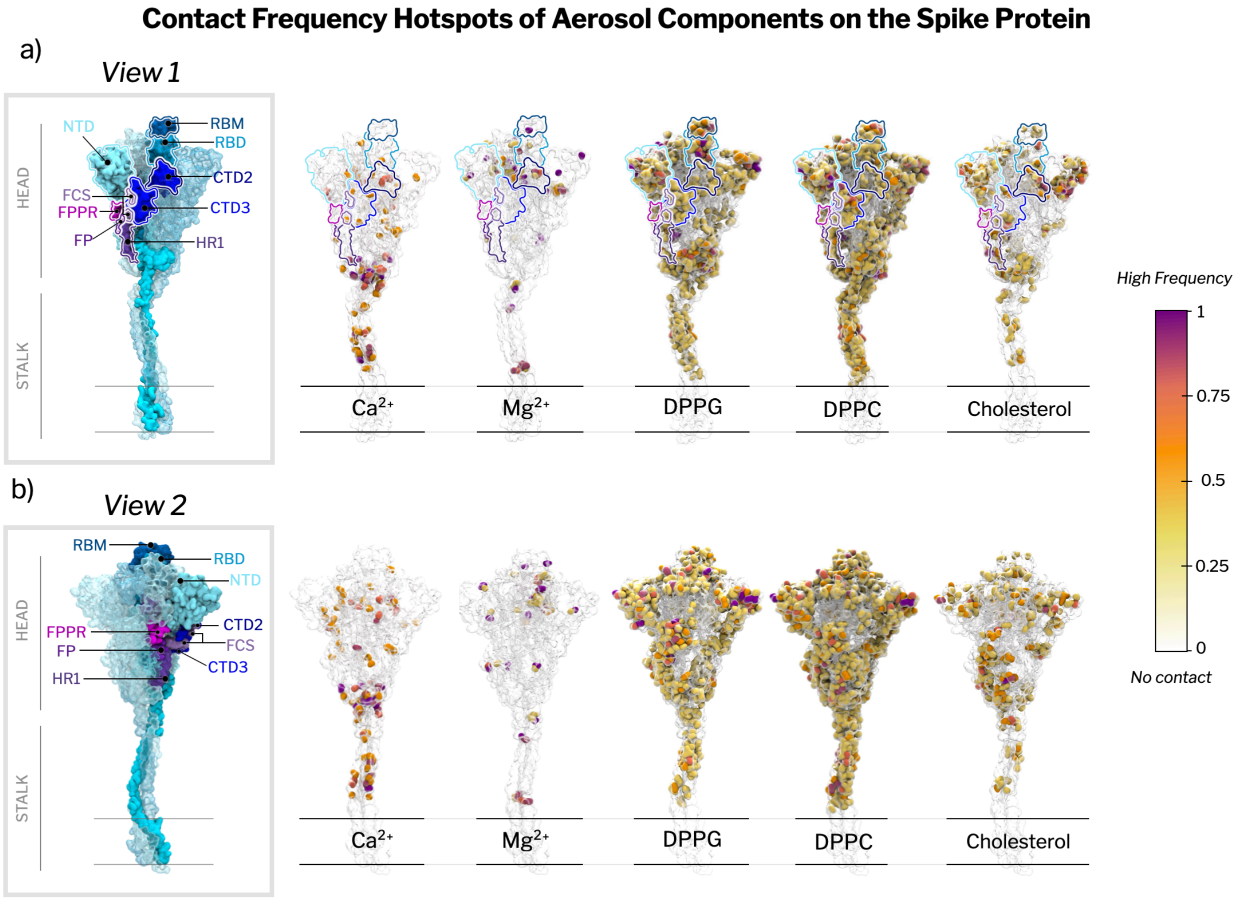


**Figure S18. Contact-frequency hotspot maps on the spike.** (a–b) Residue-level contact frequencies for major aerosol components (DPPG, DPPC, cholesterol, $Ca^{2+}$, $Mg^{2+}$) mapped onto the spike surface (plasma color scale, low (yellow) to high frequency (purple)). Panel **(a)** shows **View 1**, and panel **(b)** shows **View 2** (rotated relative to (a)) to highlight hotspots that are occluded in a single orientation. The leftmost spike in each row indicates major structural regions (head and stalk) and annotated domains (e.g., NTD/RBD/RBM, CTD2/CTD3, FCS/FPPR/FP/HR1); outlines trace these regions on the corresponding hotspot maps for reference.

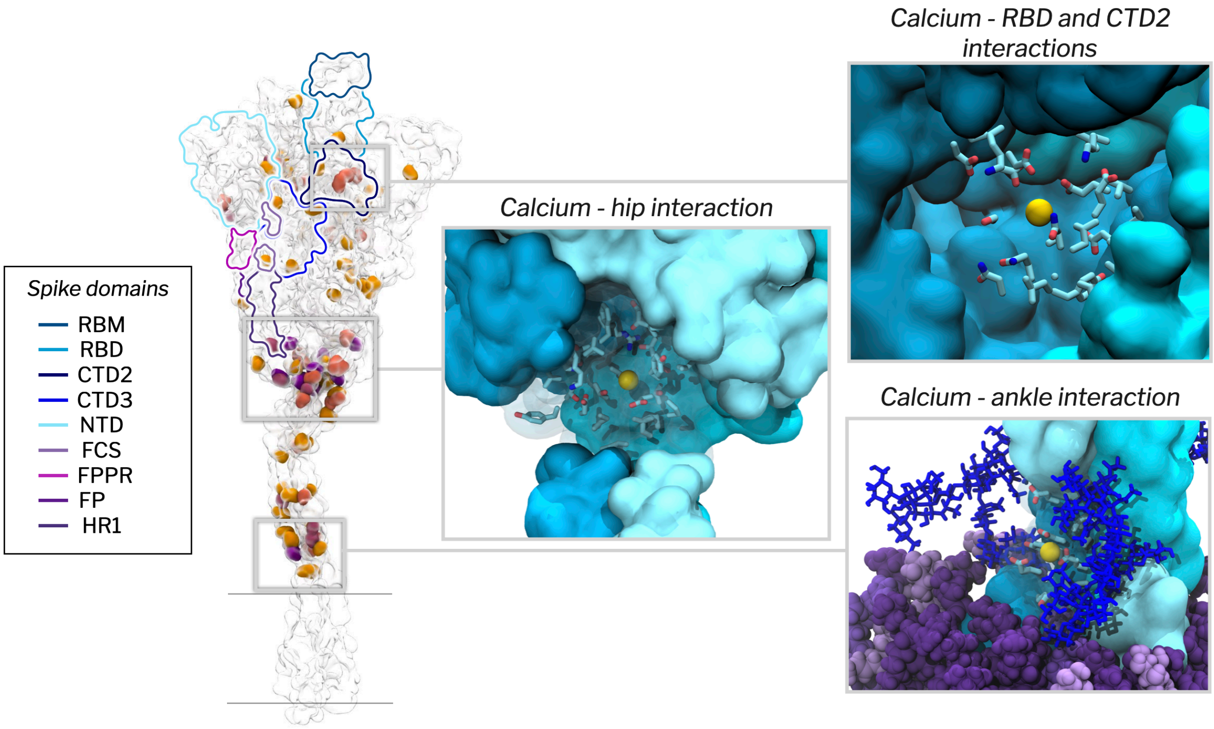


**Figure S19. Site-specific calcium enrichment and coordination environments along the spike protein.** Calcium ions display preferential accumulation at distinct structural regions of the spike, including the stalk–head junction ("hip"), the knee and ankle regions, and internal pockets within the spike head. The left panel shows the calcium contact-frequency hotspot map mapped onto the spike surface, with domains indicated. Zoomed views illustrate persistent calcium coordination at the hip (center) and ankle (bottom right), where ions are stabilized through interactions with acidic residues and nearby glycans (shown in dark blue). An additional internal pocket within the head region (top right) shows calcium occupancy coordinated by polar and charged side chains, consistent with localized stabilization. This internal pocket is located between the RBD and CTD2 domains.

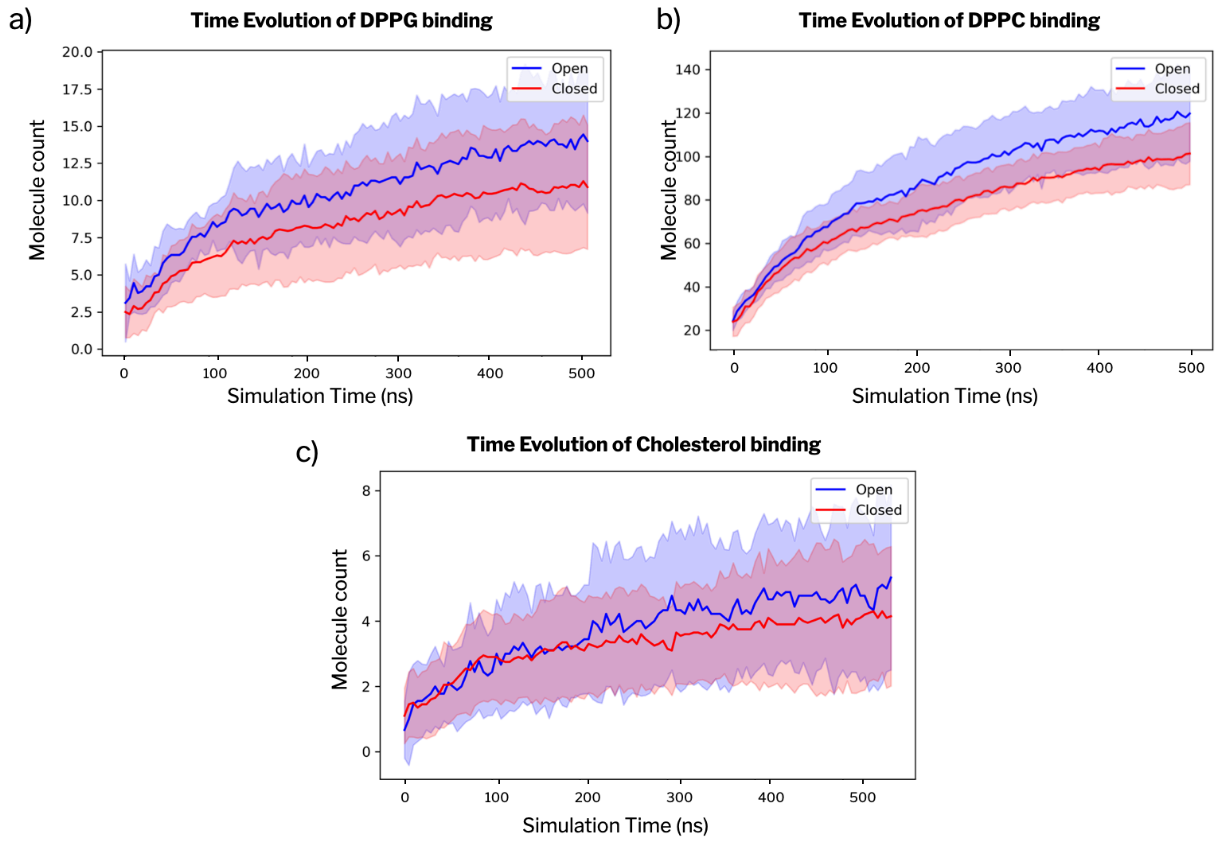


**Figure S20. Time evolution of aerosol lipid recruitment to open and closed spike conformations.** Time evolution of the number of aerosol lipid molecules in contact with spike proteins adopting open (blue) or closed (red) RBD conformations is shown for (a) DPPG, (b) DPPC, and (c) cholesterol. Solid lines represent the mean number of bound lipid molecules averaged over all spikes in each conformational state, while shaded regions indicate the corresponding variability across the ensemble. Across all lipid species, spikes with an open RBD conformation consistently recruit a larger number of lipid molecules over the course of the simulation, resulting in approximately 30% higher lipid association compared to closed conformations at long times. In contrast, analogous analyses for ionic species show no systematic dependence on spike conformation (see Figure S19).

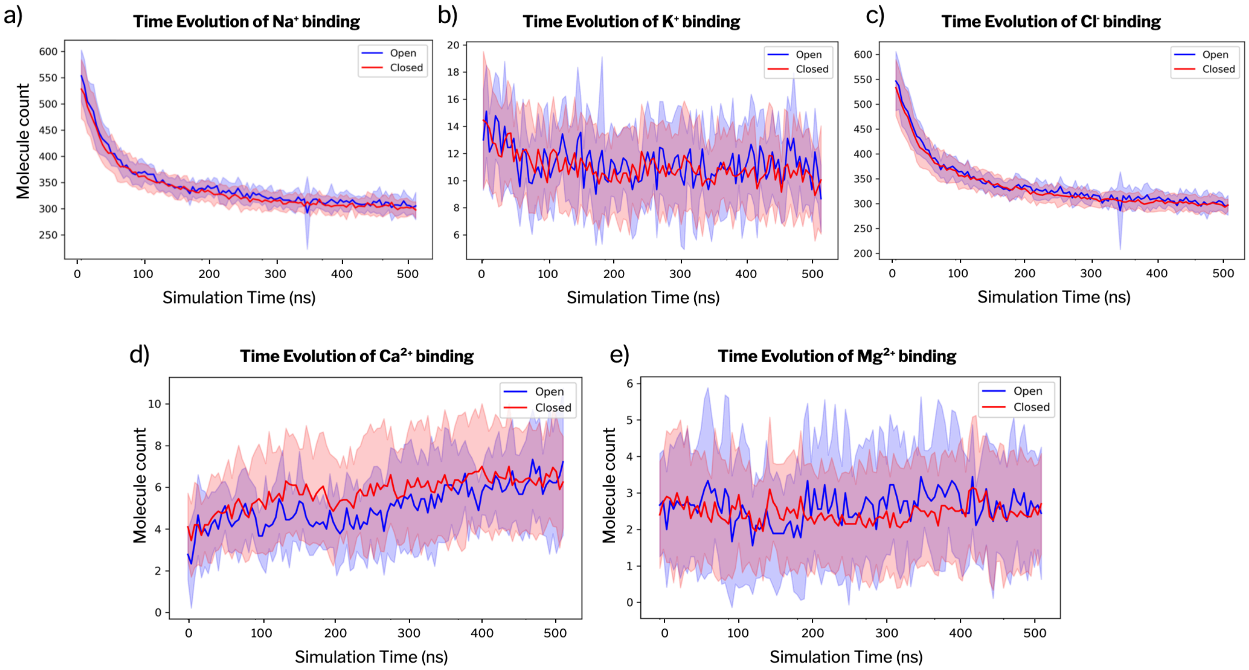


**Figure S21. Conformation-independent association of ions with the spike protein.** Time evolution of the number of ions in contact with spike proteins adopting open (blue) or closed (red) RBD conformations is shown for (a) $Na^+$, (b) $K^+$, (c) $Cl^-$, (d) $Ca^{2+}$, and (e) $Mg^{2+}$. Solid lines represent the mean number of ions associated with spikes in each conformational state, averaged over all spikes, while shaded regions indicate the corresponding variability across the ensemble. For all ionic species, the time-dependent association profiles for open and closed conformations largely overlap within statistical uncertainty, indicating that ion association with the spike surface is essentially independent of RBD conformation. This behavior contrasts with the conformation-dependent recruitment observed for aerosol lipids (Figure S20).

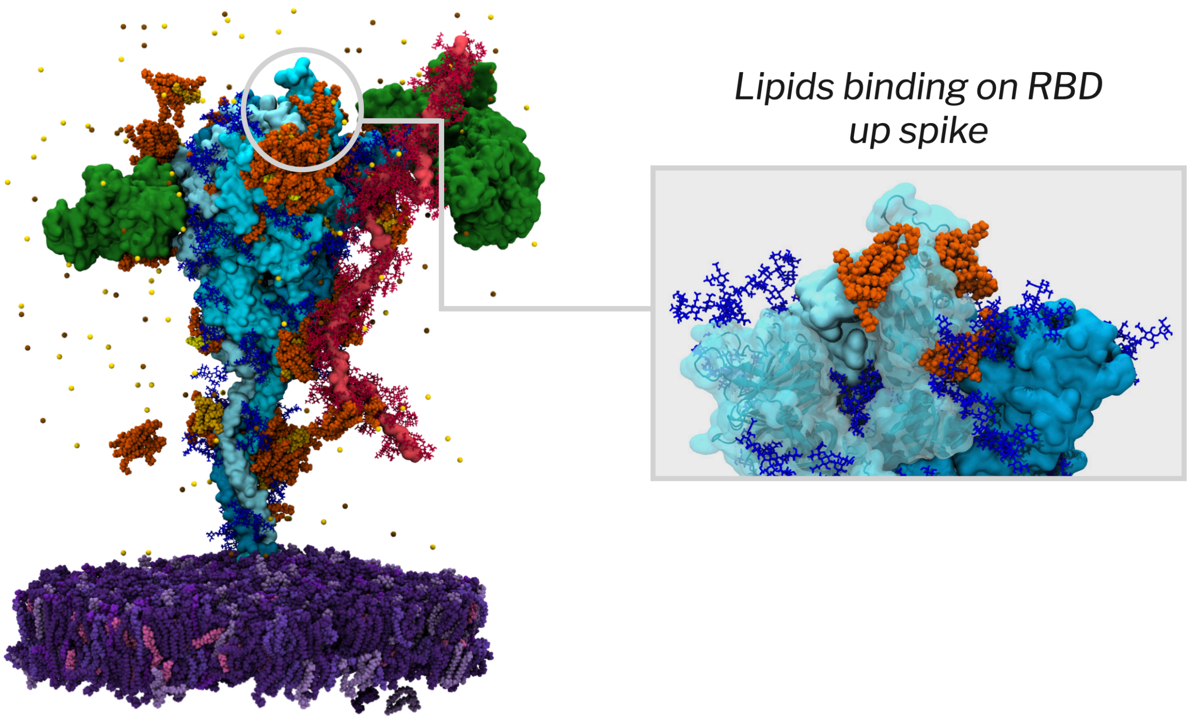


**Figure S22. Lipid recruitment at the RBD-up spike.** Representative snapshot of a spike in the open (RBD-up) conformation embedded in the viral membrane. Respiratory lipids accumulate preferentially around the exposed receptor-binding domain (RBD). Quantitative analysis shows that this lipid belt increases by ~30% relative to the closed conformation, effectively enveloping the exposed RBM and reshaping the spike microenvironment (Figure S20).

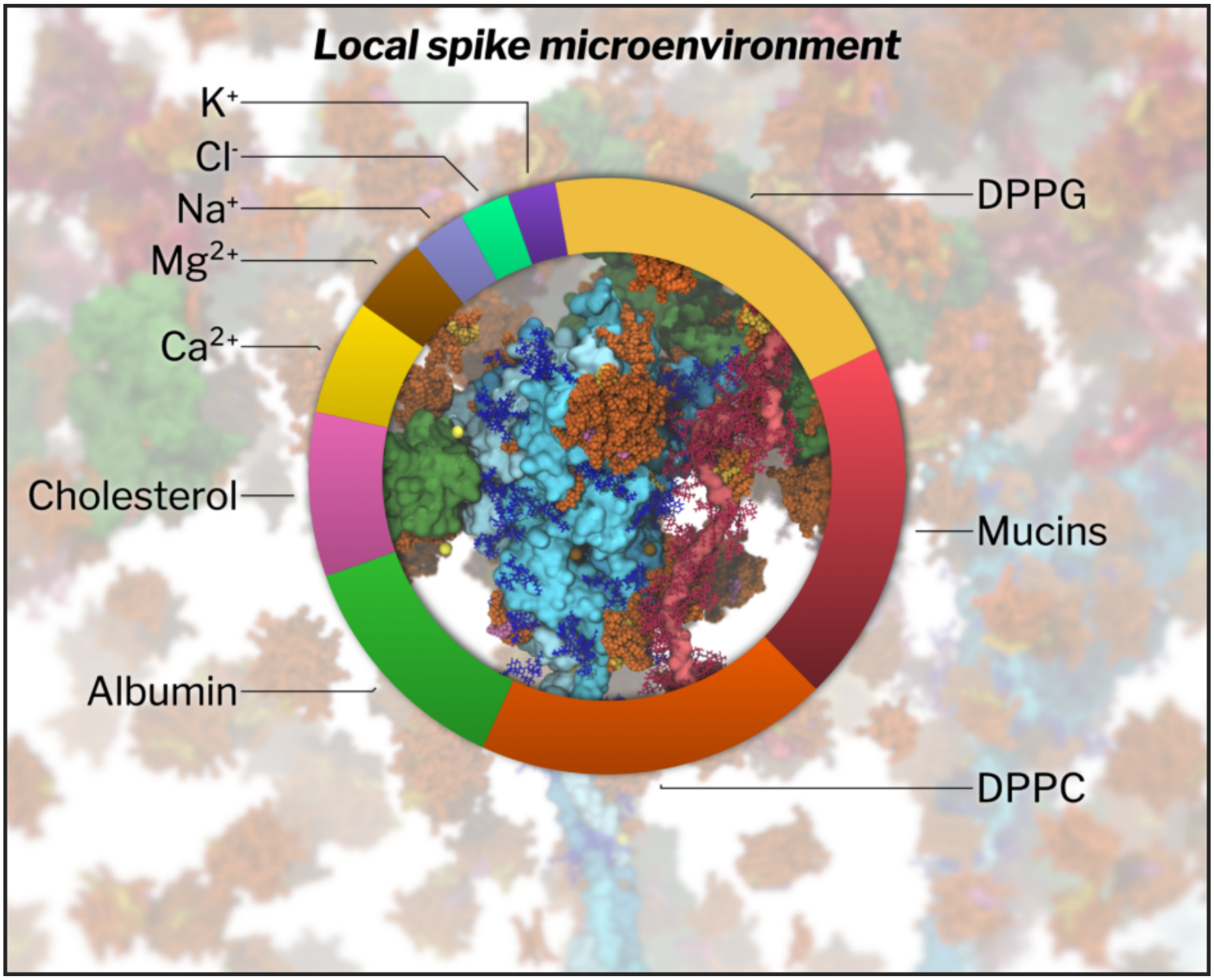


**Figure S23. Persistence interactions of molecules around the spike in the respiratory aerosol.** The relative contributions of aerosol components to persistent first-shell contacts (6 Å) of the spike over the 500 ns MD trajectory are shown. Pie segments report the fraction of cumulative contact instances attributable to each component class (DPPC, DPPG, CHL, mucins, albumin, and ions). Preference indices are reported in Table S3.

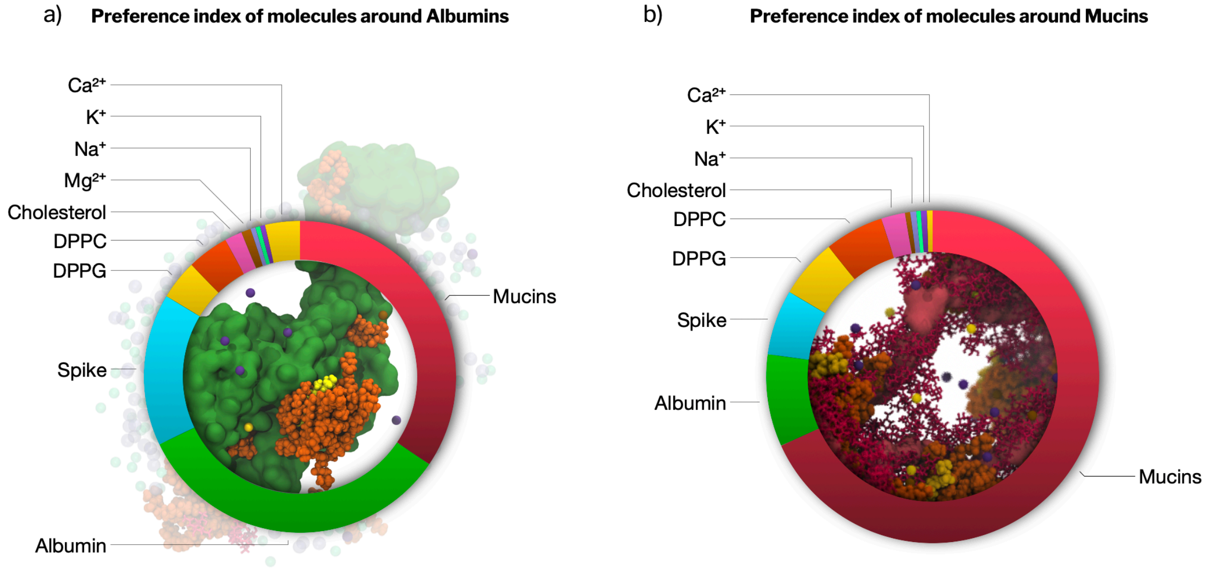


**Figure S24. Preference index of aerosol components around albumin and mucins.** The relative contributions of aerosol components to persistent first-shell contacts (within 6 Å) around (a) human serum albumin and (b) mucins are shown, computed over the 500 ns MD trajectory. Ring segments report the fraction of cumulative contact instances attributable to each component class, including spike protein, mucins, albumin, surfactant lipids (DPPC, DPPG, cholesterol), and ions. Albumin exhibits a mixed interaction environment, with substantial contributions from mucins, spike protein, and surfactant lipids, whereas mucins are predominantly surrounded by other mucin chains, reflecting their tendency to form extended, self-associated networks that dominate the local aerosol microenvironment. Preference indices are reported in Tables S4 and S5.

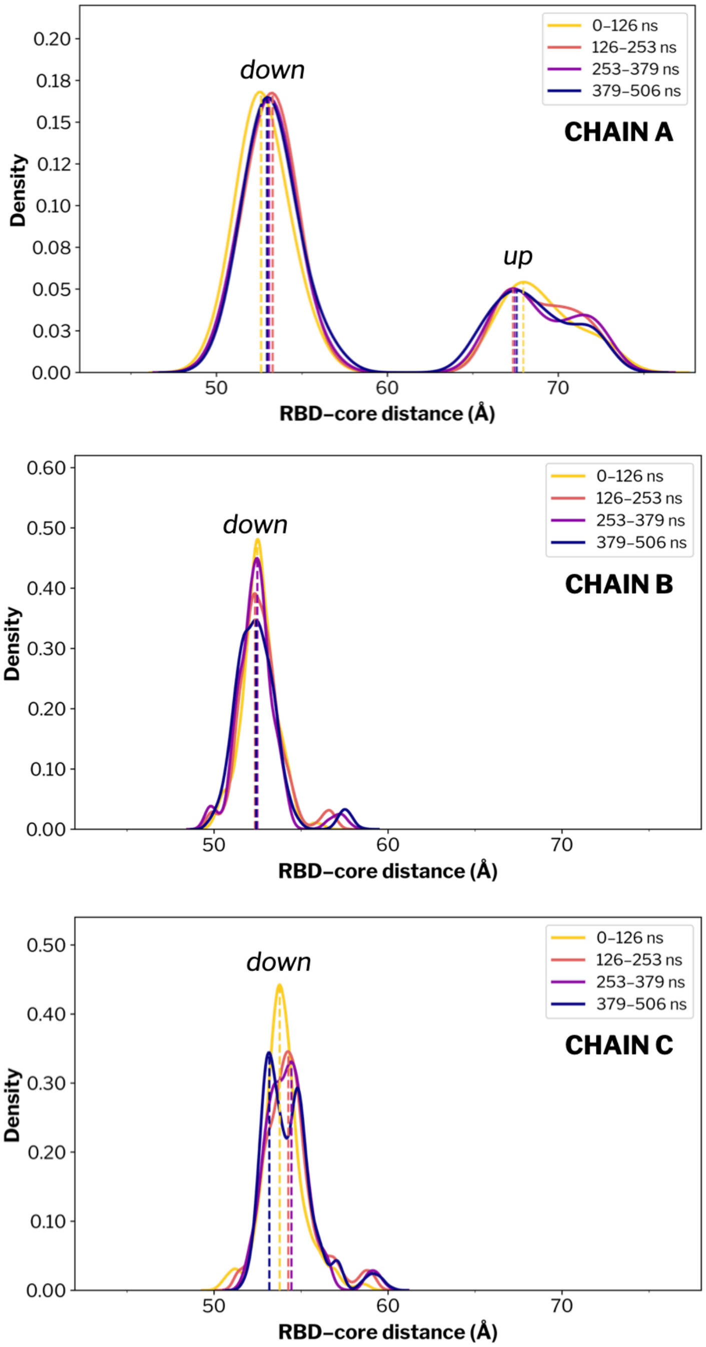


**Figure S25. RBD–core distance.** Kernel density estimates of the RBD–core distance computed across all 29 spikes for chain A (top), chain B (middle), and chain C (bottom). For each protomer, the trajectory was divided into four equal time windows of 632 frames each (~126 ns), shown from early (yellow) to late simulation time (violet), to assess temporal convergence and potential drift in the distance distributions. Peaks corresponding to RBD 'up' or 'down' are labeled accordingly.

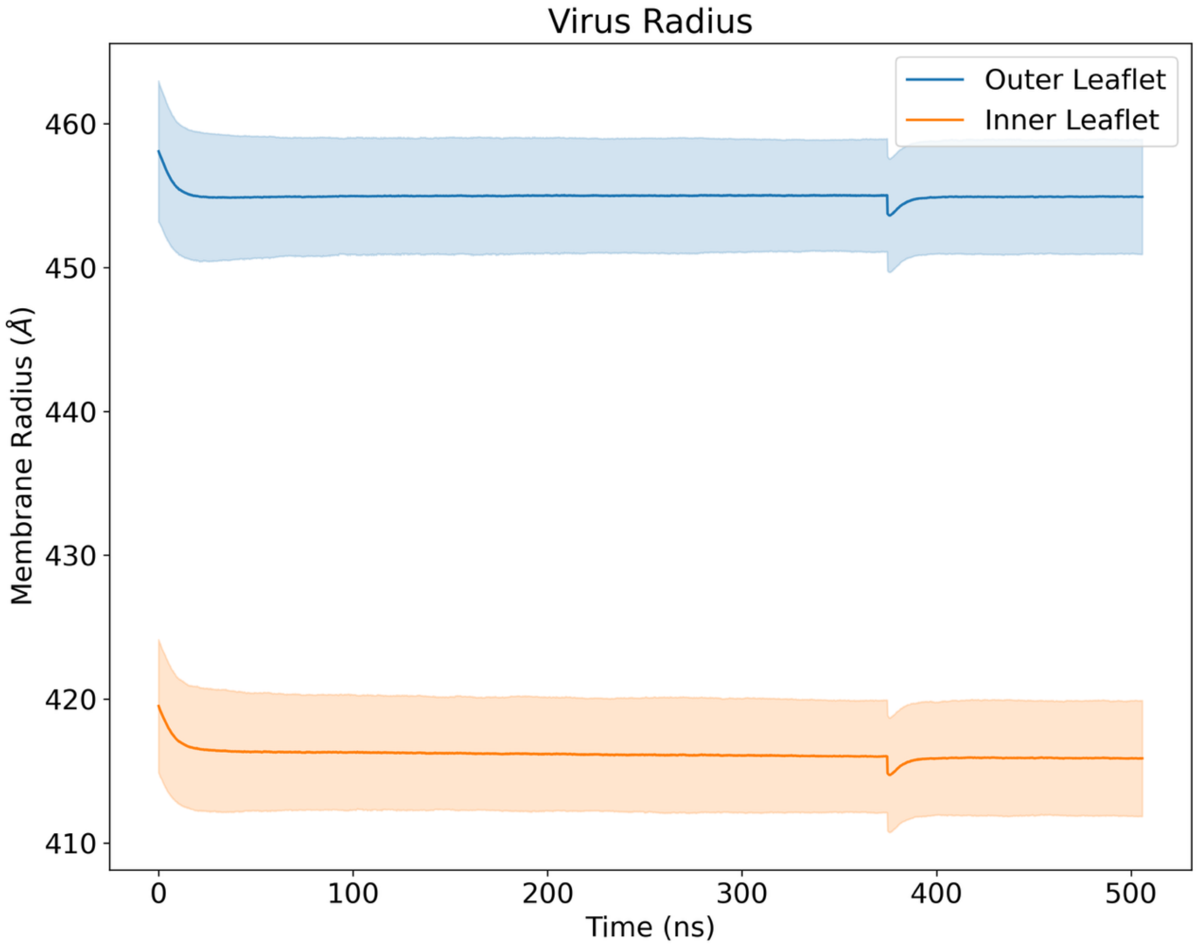


**Figure S26. Virus Membrane Radius.** The membrane radius data shows a slight shrinking at the very start of the simulation, likely due to the water density inside the virion settling causing the virus to shrink. After ~10ns the membrane remains quite stable in terms of overall diameter. There is also a noticeable dip at ~380ns after patching the membrane pore which quickly settled at a near identical radius. The outer edge of the viral membrane averages out at just around 90 nm.

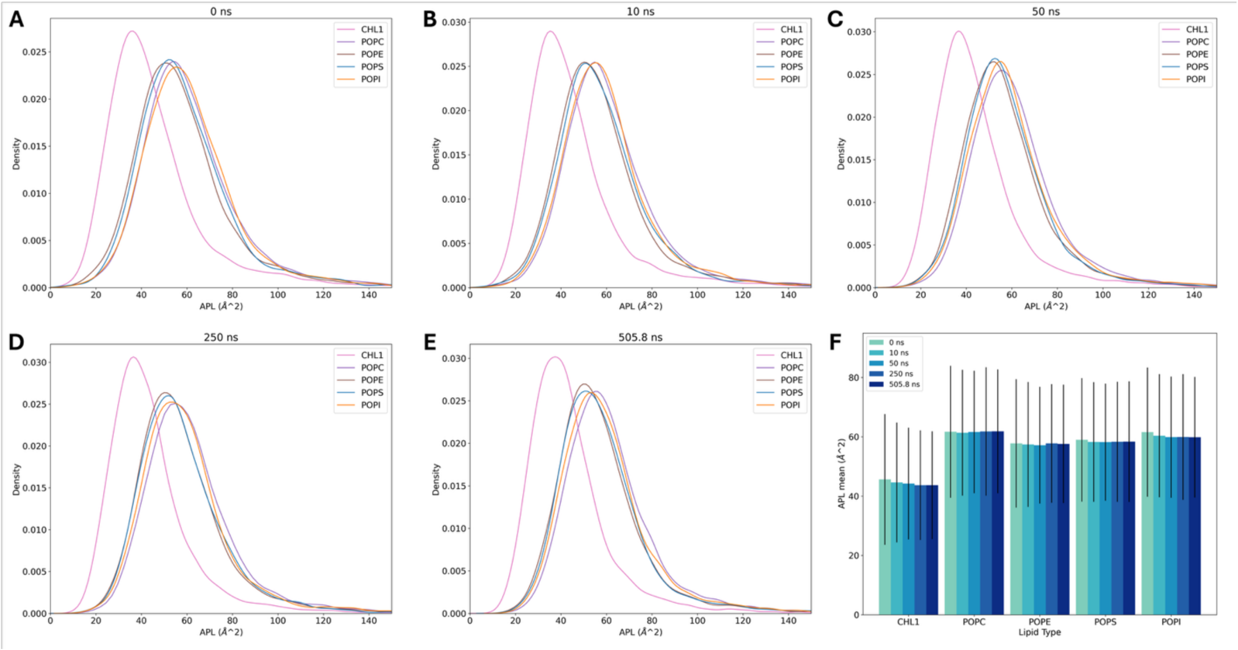


**Figure S27. Virus Membrane Data.** Area per lipid data for the first and last frames of the viral membrane. There is a minor shift to a more condensed lipid structure that follows the slight overall shrinking the virus had over the simulation.

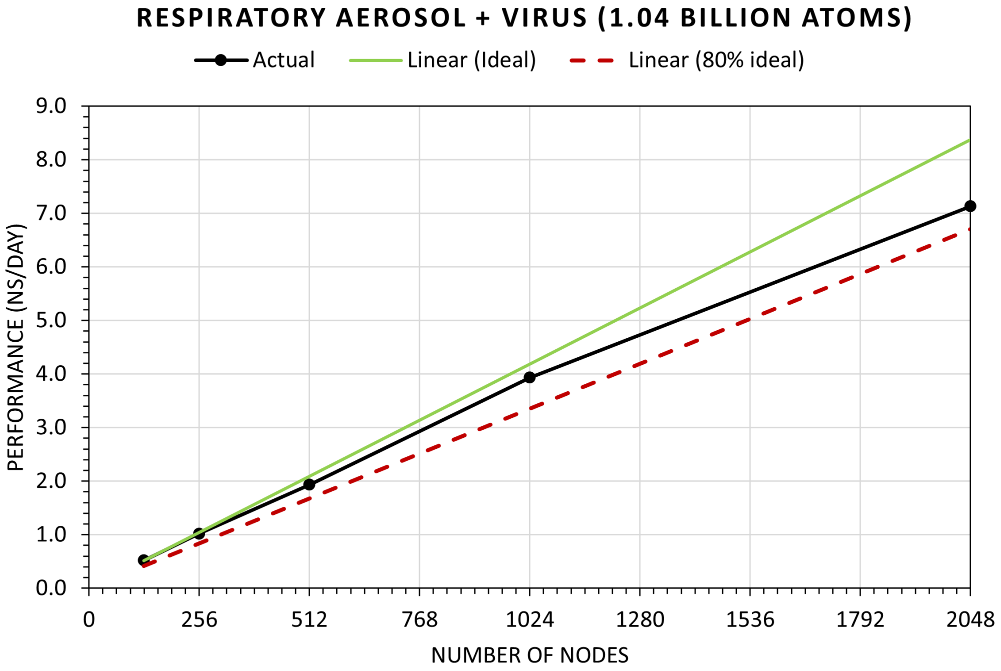

**Figure S28. System simulation scaling on TACC Frontera using Memory Optimized NAMD2.14 with a 4fs timestep and 12Å cutoff.**

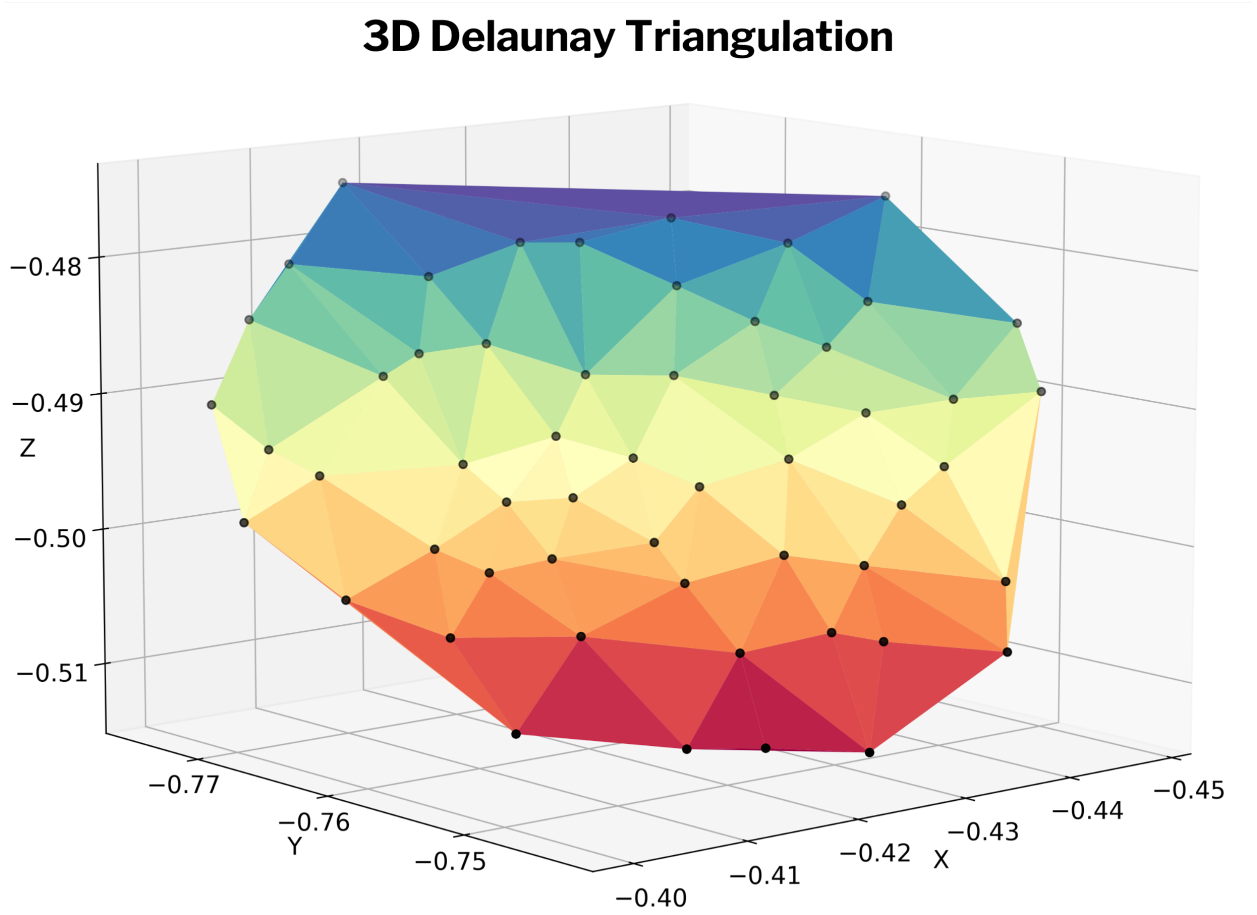


**Figure S29. Example surface lipid cluster area estimation through 3D Delaunay triangulation of the headgroup coordinates.** Each triangle area between lipid head groups is colored differently for visualization purposes.

#### *3. Supplementary Tables*

| Components | surrogate (mg/ml) | 6x surrogate | actual | % of 6x |
|---|---|---|---|---|
| **ALB** | 8.8 | 52.8 | 5.80 | 10.98% |
| **DPPG** | 0.5 | 3.0 | 2.88 | 96.00% |
| **DPPC** | 4.8 | 28.8 | 27.82 | 96.59% |
| **CHL1** | 0.1 | 0.6 | 0.58 | 96.67% |
| **PEP** | 3.0 | 18.0 | 9.64 | 53.56% |
| **HBSS** | **surrogate (mM)** | **6x surrogate** | **actual** | **% of 6x** |
| **$Ca^{2+}$** | 0.05 | 0.30 | 0.31 | 103.12% |
| **$Mg^{2+}$** | 0.02 | 0.12 | 0.11 | 97.13% |
| **$K^{+}$** | 0.23 | 1.36 | 1.22 | 89.31% |
| **$Na^{+}$** | 3.25 | 19.50 | 18.46 | 94.65% |
| **$Cl^{-}$** | 5.09 | 30.53 | 30.09 | 98.58% |

**Table S1. Quantitative composition of the RAV system relative to surrogate targets.** Surrogate lung fluid concentrations (mg $mL^{-1}$), theoretical 6× scaled concentrations final simulated ("actual") concentrations, and percent recovery relative to 6× target. Percent recovery was calculated as (actual concentration / target concentration) × 100. Simulation component labels correspond to: ALB (human serum albumin), MUC (respiratory mucins), DPPC (dipalmitoylphosphatidylcholine), DPPG (dipalmitoylphosphatidylglycerol), CHOL (cholesterol). "HBSS" refers to Hank's Balanced Salt Solution, the ionic base formulation from which the $Na^{+}$, $K^{+}$, $Ca^{2+}$, $Mg^{2+}$, and $Cl^{-}$ concentrations were derived. The final RAV composition closely matches a 6× concentration of the surrogate formulation. "Actual" reported concentrations correspond to the final respiratory fluid composition after scaling to account for aerosol evaporation from an initial ~750 nm droplet to a ~270 nm particle. Mucin content was modestly down-sampled to maintain tractable system size while preserving relative volume fraction and crowding behavior.

| | Full Dataset | Mucin Dataset |
|---|---|---|
| **Water** | 0.46 | 0.43 |
| **Resp. Lipid** | 0.08 | 0.05 |
| **Memb. Lipid** | -0.31 | -0.20 |
| **Albumin** | -0.08 | -0.19 |
| **Spike** | -0.15 | -0.17 |
| **Mucin** | -0.07 | -0.20 |
| **Divalent Cations** | 0.17 | 0.11 |
| **Ions** | 0.35 | 0.46 |

**Table S2. Pearson correlation between the calculated diffusion coefficient for a voxel and the mass of various component groups.** The full dataset only excludes regions that had too few waters to accurately estimate the diffusion whereas the mucin dataset excludes regions that had fewer than 1000 amu of mucin atoms.

| Component | Mean contacts (<6 Å) | SD | Total available in aerosol | NBR | Preference Index (PI) |
|---|---|---|---|---|---|
| M protein | 1,34 | 1,01 | 720 | $1.86 \times 10^{-3}$ | 41,48 |
| DPPG | 11,62 | 4,32 | 24807 | $4.68 \times 10^{-4}$ | 10,45 |
| DPPC | 102,44 | 16,27 | 236055 | $4.34 \times 10^{-4}$ | 9,68 |
| Mucins | 0,13 | 0,32 | 345 | $3.83 \times 10^{-4}$ | 8,53 |
| Albumin | 0,16 | 0,50 | 550 | $2.88 \times 10^{-4}$ | 6,41 |
| Cholesterol | 4,28 | 2,13 | 21213 | $2.02 \times 10^{-4}$ | 4,50 |
| $Ca^{2+}$ | 6,37 | 1,92 | 44827 | $1.42 \times 10^{-4}$ | 3,17 |
| $Mg^{2+}$ | 2,61 | 1,16 | 27281 | $9.58 \times 10^{-5}$ | 2,14 |
| $Na^{+}$ | 306,46 | 10,98 | 4627417 | $6.62 \times 10^{-5}$ | 1,48 |
| $Cl^{-}$ | 302,04 | 11,43 | 4885636 | $6.18 \times 10^{-5}$ | 1,38 |
| $K^{+}$ | 10,67 | 1,08 | 179254 | $5.95 \times 10^{-5}$ | 1,33 |
| E protein | 0 | 0 | 20 | 0 | 0 |

**Table S3. Preference index analysis for the 6 Å interaction shell around the spike protein.** For each component, the time-averaged number of molecules within 6 Å of the spike, total abundance in the aerosol, normalized binding ratio (NBR), and preference index (PI) are reported. Preference indices were calculated using a single pooled global mean NBR ($4.48 \times 10^{-5}$), computed across all reference interactomes (spike, albumin, and mucins), and applied as a fixed normalization constant. Spike–spike interactions were excluded due to membrane anchoring. Values of PI > 1 indicate enrichment relative to the global aerosol average, whereas PI < 1 indicates depletion.

| Component | Mean contacts (<6 Å) | SD | Total available in aerosol | NBR | Preference Index (PI) |
|---|---|---|---|---|---|
| Mucins | 0,22 | 0,45 | 345 | $6.52 \times 10^{-4}$ | 14,54 |
| Albumin | 0,33 | 0,59 | 550 | $6.09 \times 10^{-4}$ | 13,58 |
| Spike | 0,01 | 0,08 | 29 | $2.88 \times 10^{-4}$ | 6,41 |
| DPPC | 18,59 | 7,58 | 236055 | $7.87 \times 10^{-5}$ | 1,76 |
| DPPG | 1,86 | 1,43 | 24807 | $7.50 \times 10^{-5}$ | 1,67 |
| $Ca^{2+}$ | 2,97 | 1,15 | 44827 | $6.61 \times 10^{-5}$ | 1,48 |
| Cholesterol | 0,70 | 0,79 | 21213 | $3.31 \times 10^{-5}$ | 0,74 |
| $Mg^{2+}$ | 0,49 | 0,34 | 27281 | $1.78 \times 10^{-5}$ | 0,40 |
| $Na^{+}$ | 44,58 | 2,55 | 4627417 | $9.63 \times 10^{-6}$ | 0,21 |
| $K^{+}$ | 1,69 | 0,32 | 179254 | $9.45 \times 10^{-6}$ | 0,21 |
| $Cl^{-}$ | 44,53 | 3,01 | 4885636 | $9.11 \times 10^{-6}$ | 0,20 |
| M protein | 0,00 | 0,01 | 720 | $3.48 \times 10^{-7}$ | 0,01 |
| E protein | 0 | 0 | 20 | 0 | 0 |

**Table S4. Preference index analysis for the 6 Å interaction shell around albumin.** For each component, the time-averaged number of molecules within 6 Å of albumin, total aerosol abundance, normalized binding ratio (NBR), and preference index (PI) are shown. The NBR is defined as the ratio between the mean number of contacting molecules and the total number of that component present in the aerosol. Preference indices were normalized using the same pooled global mean NBR ($4.48 \times 10^{-5}$) used for the spike and mucin analyses, enabling direct comparison across reference surfaces. PI values greater than unity indicate preferential enrichment.

| Component | Mean contacts (<6 Å) | SD | Total available in aerosol | NBR | Preference Index (PI) |
|---|---|---|---|---|---|
| Mucins | 1,58 | 1,23 | 345 | $4.57 \times 10^{-3}$ | 102,00 |
| Albumin | 0,36 | 0,57 | 550 | $6.51 \times 10^{-4}$ | 14,53 |
| DPPC | 93,73 | 17,38 | 236055 | $3.97 \times 10^{-4}$ | 8,86 |
| DPPG | 9,83 | 3,61 | 24807 | $3.96 \times 10^{-4}$ | 8,84 |
| Spike | 0,01 | 0,10 | 29 | $3.83 \times 10^{-4}$ | 8,53 |
| Cholesterol | 3,39 | 1,84 | 21213 | $1.60 \times 10^{-4}$ | 3,56 |
| $Na^+$ | 190,19 | 16,56 | 4627417 | $4.11 \times 10^{-5}$ | 0,92 |
| $K^+$ | 7,32 | 0,91 | 179254 | $4.08 \times 10^{-5}$ | 0,91 |
| $Ca^{2+}$ | 1,73 | 0,47 | 44827 | $3.86 \times 10^{-5}$ | 0,86 |
| $Mg^{2+}$ | 0,95 | 0,30 | 27281 | $3.49 \times 10^{-5}$ | 0,78 |
| $Cl^-$ | 145,68 | 14,49 | 4885636 | $2.98 \times 10^{-5}$ | 0,67 |
| M protein | 0,00 | 0,07 | 720 | $4.86 \times 10^{-6}$ | 0,11 |
| E protein | 0 | 0 | 20 | 0 | 0 |

**Table S5. Preference index analysis for the 6 Å interaction shell around mucins.** For each component the mean contact count (<6 Å), total aerosol abundance, normalized binding ratio (NBR), and preference index (PI) are reported. Preference indices were calculated using the same pooled global mean NBR ($4.48 \times 10^{-5}$) applied across all reference systems. PI values above 1 indicate enrichment relative to the overall aerosol molecular ensemble.

| Component | Full RAV | Dry RAV |
|---|---|---|
| **Water** | **89.4%** | **-** |
| **Organics** | **5.8%** | **54.6%** |
| Virion | 1.3% | 12.3% |
| Resp. Fluid | 4.5% | 42.3% |
| Mucin | 0.9% | 8.7% |
| Albumin | 0.6% | 5.2% |
| DPPC | 2.7% | 25.2% |
| DPPG | 0.3% | 2.6% |
| Chol. | 0.1% | 0.5% |
| **Ions** | **4.8%** | **45.4%** |
| $Na^+$ | 0.0% | 0.3% |
| $Cl^-$ | 2.9% | 27.2% |
| $Ca^{2+}$ | 1.8% | 16.7% |
| $Mg^{2+}$ | 0.0% | 0.1% |
| $K^+$ | 0.1% | 1.1% |

Table S6. Categorized mass percent breakdown for the RAV, both with water included and without.

***4. Supplementary References***


Blender Online Community. (2020). *Blender—A 3D modelling and rendering package* (Version 2.79b). Blender Foundation. http://www.blender.org

Casalino, L., Gaieb, Z., Goldsmith, J. A., Hjorth, C. K., Dommer, A. C., Harbison, A. M., Fogarty, C. A., Barros, E. P., Taylor, B. C., McLellan, J. S., Fadda, E., & Amaro, R. E. (2020). Beyond Shielding: The Roles of Glycans in the SARS-CoV-2 Spike Protein. *ACS Central Science*, *6*(10), 1722–1734. https://doi.org/10.1021/acscentsci.0c01056

Corradi, V., Mendez-Villuendas, E., Ingólfsson, H. I., Gu, R.-X., Siuda, I., Melo, M. N., Moussatova, A., DeGagné, L. J., Sejdiu, B. I., Singh, G., Wassenaar, T. A., Delgado Magnero, K., Marrink, S. J., & Tieleman, D. P. (2018). Lipid–Protein Interactions Are Unique Fingerprints for Membrane Proteins. *ACS Central Science*, *4*(6), 709–717. https://doi.org/10.1021/acscentsci.8b00143

Dommer, A., Casalino, L., Kearns, F., Rosenfeld, M., Wauer, N., Ahn, S.-H., Russo, J., Oliveira, S., Morris, C., Bogetti, A., Trifan, A., Brace, A., Sztain, T., Clyde, A., Ma, H., Chennubhotla, C., Lee, H., Turilli, M., Khalid, S., … Amaro, R. E. (2023). #COVIDisAirborne: AI-enabled multiscale computational microscopy of delta SARS-CoV-2 in a respiratory aerosol. *The International Journal of High Performance Computing Applications*, *37*(1), 28–44. https://doi.org/10.1177/10943420221128233

Durrant, J. D., & Amaro, R. E. (2014). LipidWrapper: An Algorithm for Generating Large-Scale Membrane Models of Arbitrary Geometry. *PLoS Computational Biology*, *10*(7). https://doi.org/10.1371/journal.pcbi.1003720

Hopkins, C. W., Le Grand, S., Walker, R. C., & Roitberg, A. E. (2015). Long-Time-Step Molecular Dynamics through Hydrogen Mass Repartitioning. *Journal of Chemical Theory and Computation*, *11*(4), 1864–1874. https://doi.org/10.1021/ct5010406

Humphrey, W., Dalke, A., & Schulten, K. (1996). VMD: Visual Molecular Dynamics. *Journal of Molecular Graphics*, *14*(February), 33–38. https://doi.org/10.1016/0263-7855(96)00018-5

Jo, S., Kim, T., Iyer, V. G., & Im, W. (2008). CHARMM-GUI: A web-based graphical user interface for CHARMM. *Journal of Computational Chemistry*, *29*(11), 1859–1865. https://doi.org/10.1002/jcc.20945

Ke, Z., Oton, J., Qu, K., Cortese, M., Zila, V., McKeane, L., Nakane, T., Zivanov, J., Neufeldt, C. J., Cerikan, B., Lu, J. M., Peukes, J., Xiong, X., Kräusslich, H.-G., Scheres, S. H. W., Bartenschlager, R., & Briggs, J. A. G. (2020). Structures and distributions of SARS-CoV-2 spike proteins on intact virions. *Nature*, *588*(7838), 498–502. https://doi.org/10.1038/s41586-020-2665-2

Mandala, V. S., McKay, M. J., Shcherbakov, A. A., Dregni, A. J., Kolocouris, A., & Hong, M. (2020). Structure and drug binding of the SARS-CoV-2 envelope protein transmembrane domain in lipid bilayers. *Nature Structural and Molecular Biology*, *27*(12), 1202–1208.

Michaud-Agrawal, N., Denning, E. J., Woolf, T. B., & Beckstein, O. (2011). MDAnalysis: A toolkit for the analysis of molecular dynamics simulations. *Journal of Computational Chemistry*, *32*(10), 2319–2327. https://doi.org/10.1002/jcc.21787

Shukla, D., Shinde, C., & Trout, B. L. (2009). Molecular Computations of Preferential Interaction Coefficients of Proteins. *The Journal of Physical Chemistry B*, *113*(37), 12546–12554. https://doi.org/10.1021/jp810949t

Surya, W., Li, Y., & Torres, J. (2018). Structural model of the SARS coronavirus E channel in LMPG micelles. *Biochimica et Biophysica Acta (BBA) - Biomembranes*, *1860*(6), 1309–1317. https://doi.org/10.1016/j.bbamem.2018.02.017

Sztain, T., Ahn, S.-H., Bogetti, A. T., Casalino, L., Goldsmith, J. A., Seitz, E., McCool, R. S., Kearns, F. L., Acosta-Reyes, F., Maji, S., Mashayekhi, G., McCammon, J. A., Ourmazd, A., Frank, J., McLellan, J. S., Chong, L. T., & Amaro, R. E. (2021). A glycan gate controls opening of the SARS-CoV-2 spike protein. *Nature Chemistry 2021 13:10*, *13*(10), 963–968. https://doi.org/10.1038/s41557-021-00758-3

Vejerano, E. P., & Marr, L. C. (2018). Physico-chemical characteristics of evaporating respiratory fluid droplets. *Journal of the Royal Society Interface*, *15*(139), 1–10. https://doi.org/10.1098/rsif.2017.0939

Walker, J. S., Archer, J., Gregson, F. K. A., Michel, S. E. S., Bzdek, B. R., & Reid, J. P. (2021). Accurate Representations of the Microphysical Processes Occurring during the Transport of Exhaled Aerosols and Droplets. *ACS Central Science*, *7*(1), 200–209. https://doi.org/10.1021/acscentsci.0c01522

Wexler, A., & Greenspan, L. (1971). *I Vapor Pressure Equation for Water in the Range 0 to 100°C. Vol. 75A*(No. 3).
